# Polyphosphate metabolism genes are associated with hydroxyapatite biomineralization capacity in a novel hydrogenotrophic methanogen *Methanobacterium* AMB_03

**DOI:** 10.64898/2026.11.06.764883

**Authors:** F. A. Robin-Champigneul, A. Singh, S. Sharma, O. Tkachenko, J. Thyr, S. Sidiropoulos, A. Schnürer, T. Zajkowski, T. Budnyak, A. Neubeck

**Affiliations:** Department of Earth Science - Centre for Deep Sea Research, University of Bergen, Norway; Department of Microbiology, Swedish University of Agricultural Sciences, Sweden; Blue Marble Space Institute of Science, Seattle, WA, 98104, USA; Department of Earth Sciences, Uppsala University, Sweden; Department of Material Science and Engineering: solid state physics, Uppsala University, Sweden; AGH University, Faculty of Space Technologies, 30-059 Kraków, Poland; Wallenberg Initiative Materials Science for Sustainability, Department of Earth Sciences, Uppsala University, Villavägen 16, 752 36 Uppsala, Sweden

**Keywords:** polyP, hydrogenotrophic methanogen, hydroxyapatite, biomineralization, phosphate genomes

## Abstract

Methanogenic archaea play crucial roles in sedimentary phosphorus cycling, but their capacity for phosphate biomineralization remains poorly understood. Here we demonstrate that *Methanobacterium* AMB_03, isolated from a serpentinizing environment, mediates extracellular precipitation of carbonate-substituted hydroxyapatite (HAp) under phosphate-enriched conditions. Comparative genomic analysis of AMB_03 versus non-mineralizing strains (*Methanobacterium oryzae*, *Methanoculleus bourgensis* MAB1) shows that AMB_03 encodes polyphosphate kinase 1 (*ppk1*), enabling polyphosphate synthesis from ATP. In contrast, *M. oryzae* and MAB1 completely lack *ppk1*, preventing polyphosphate accumulation. Phylogenomic analysis of 320 *Methanobacteriota* genomes further shows that complete polyP cycling capacity is present in only ∼25% of representatives, placing AMB_03 among a minority of methanogens with this capability. Experimental cultivation across phosphate gradients (0-6 mM) showed HAp early-onset precipitation was preferential at ≥3 mM. Precipitation was eventually observed at all phosphate concentrations (except P0) by 49 weeks and was associated with extracellular polymeric substance production and localized pH increases of up to 8.08. Raman spectroscopy confirmed carbonate-substituted HAp, with characteristic peaks at 960.5 cm^⁻¹^ (PO_4_^3-^) and 1070-1072 cm⁻¹ (CO_3_^2-^). These findings identify a strong genomic association between polyphosphate metabolism genes and biomineralization capacity, suggesting that methanogens with complete polyP cycling systems may create localized phosphate-rich environments that promote authigenic mineral formation in sedimentary systems.

## 1. Introduction

### 1.1 Methanogens and polyP

The mechanisms by which methanogens manage phosphorus under variable availability remain poorly constrained but may have been critical to nutrient cycling in early anoxic environments. Paula *et al*. (2019) demonstrated that methanogenic Archaea, including *Methanosarcina mazei*, can accumulate intracellular inorganic polyphosphate (polyP) under anaerobic conditions following inorganic phosphate starvation and subsequent re-exposure. This accumulation coincided with transcriptional upregulation of the gene *ppk1*, indicating that polyP metabolism in methanogens operates independently of oxygen and may thus represent an ancient cellular strategy for phosphate storage and regulation. Several studies have reported that polyP accumulation is widespread among methanogens, occurring not only in hydrogenotrophic species but also in those using other substrates (Rudnick et al., 1990; Scherer et al., 1983; Toso et al., 2011). Polyphosphate kinase 1 synthesizes polyphosphate by transferring terminal phosphates from ATP, while exopolyphosphatase (*ppx*) degrades polyphosphate by cleaving terminal phosphate residues. Together, *ppk1* and *ppx* constitute a synthesis-degradation cycle that, while first characterized in bacteria, is encoded by homologous genes in polyP-capable methanogens (Paula et al. 2019; section 3.6). The broader physicochemical and regulatory roles of polyP have been characterized primarily in bacteria, and are thought to be conserved given the antiquity of the molecule (Racki & Freddolino, 2025). PolyP is a negatively charged molecule that forms biomolecular condensates in bacterial cells, droplet-like structures that help organize cellular processes without membranes (Racki and Freddolino, 2025). These condensates often cluster around the nucleoid and are important for stress survival, gene regulation, and cell cycle control (Racki and Freddolino, 2025). It interacts with metals (like Mg^2+^ and Mn^2+^), proteins, and small metabolites, which together determine how DNA and RNA behave within the cell. Metals can drive the physical formation of polyP droplets and influence DNA folding. When bacteria lose the ability to make polyP, they struggle to survive stress, their chromosomes become unstable, and gene expression becomes misregulated, showing that polyP is central to bacterial organization and adaptation (Rao and Kornberg, 1996). PolyP accumulation itself is well documented also across methanogens in the archaeal domain (Rudnick et al. 1990; Scherer et al. 1983; Toso et al. 2011; Paula et al. 2019), but whether it performs the same condensate-forming role described in bacteria remains untested in archaea. Within methanogens specifically, phosphate limitation has been shown to decouple catabolism and anabolism and to reduce methanogenesis capacity (Gu et al., 2022; Paula et al., 2019). In this case, cells restrict methane production not because of limited energy substrates, but because phosphate scarcity prevents growth and triggers regulatory down-shifts in catabolic activity. In contrast, other studies have shown that hydrogenotrophic methanogens tend to be limited by their energy substrates rather than by phosphorus availability (Medvedeff et al., 2014), showing that hydrogenotrophic methanogens predominantly experience energy limitation driven by restricted H_2_/CO_2_ supply, rather than nutrient limitation imposed by low phosphate concentrations. This apparent contrast draws on studies using different methanogen taxa and experimental systems (pure-cultures and natural soil community), so it is not yet clear whether energy-vs. phosphate-limitation reflects a true taxon-or pathway-level difference, or just differences in experimental context; a related pattern in *M.* AMB_03 is discussed in Sections 3.1 and 4.3. Understanding polyphosphate metabolism in methanogens becomes particularly relevant when considering their role in sedimentary phosphorus cycling.

### 1.2 Phosphorus cycling

The oxygenation of the oceans is maintained by a fine balance between the supply of P and the burial of organic carbon. This balance is ‘fine’ because even relatively small changes in nutrient delivery or organic-matter burial can shift the system toward either well-oxygenated or O_2_-poor conditions. An increased supply of P, such as from enhanced terrestrial weathering or reduced burial under low O_2_ conditions, can stimulate primary productivity and increase the export of organic matter to depth. Microbial respiration of this accumulating organic matter consumes dissolved O_2_ in subsurface, oceanic waters. If P inputs or biological utilization become high enough, the resulting increase in O_2_ demand can expand low O_2_ zones and push parts of the ocean closer to anoxia (Watson et al., 2017). During warm periods, terrestrial weathering increases and delivers increased amounts of P to the oceans, triggering primary productivity, O_2_ consumption and oceanic anoxia (Watson et al., 2017). Several events of abrupt increase in nutrient supply to the oceans in the past have been shown to be one of the main causes for mass extinctions (Kidder and Worsley, 2010). Within these sedimentary systems, the methanogenic zone represents a critical interface where phosphorus transformations occur.

### 1.3 Marine sediment methanogen zone and phosphate formation

Sedimentary phosphorites are the largest sink of phosphorus in the rock record and influence the marine primary production, both today and through geological time. The formation of marine phosphate minerals is commonly associated with environments with high primary production together with a high organic matter burial (Mänd et al., 2018). Continental margin upwelling zones are examples of such environments where the deposition is high and where the terrigenous Fe (oxyhydr)oxides (FeOx) are the main carriers of adsorbed P (März et al., 2018). The precipitation of phosphates occurs mainly in the sediment column; the FeOx reaches below the oxic zone and starts to dissolve, releasing the P into the sediment pore waters. The fate of P is then either to reprecipitate within the sediment column or to migrate up to the water column. The reprecipitation of phosphates is especially prominent in the sulfate-methane transition zone (SMT) where dissolution of FeOx occurs. However, some of the P released into the pore waters by diagenetic processes, are also migrating downwards to the methanogenic zone, where it may precipitate and become buried over long time scales (März et al., 2018).

Authigenic apatite (commonly carbonate fluorapatite, CFA) has widely been regarded as the dominant long-term phosphorus sink in marine sediments (Delaney, 1998; Filippelli, 1997; März et al., 2018). However, increasing evidence now shows that Fe(II) phosphates such as vivianite, classically associated with freshwater systems, also form in marine environments (Berner, 1990; Bray et al., 1973; Fagel et al., 2005). Vivianite can develop where intense organic matter degradation rapidly depletes sulfate, limiting sulfide formation and allowing Fe^2+^ to accumulate and precipitate with phosphate (Egger et al., 2015; Hsu et al., 2014; Li et al., 2015; März et al., 2018). Even in the fully anoxic Black Sea, Fe(II) phosphates may form below the sulfide penetration depth (Egger et al., 2016). Similar Fe - P coupling below the SMT has been reported in deep-marine settings such as the Amazon Fan and the Zambezi deep-sea fan (Burns, 1997; März et al., 2008). Below the SMT in marine sediments, phosphate is shown not to primarily be removed by re-adsorption onto FeOx or by authigenic CFA formation, because buried FeOx loses reactivity and carbonate-poor sediments limit CFA precipitation (März et al., 2018). Instead, the main sink for phosphate appears to be the formation of finely dispersed Fe(II) phosphate minerals, such as vivianite, which form under anoxic, reducing conditions where dissolved Fe^2+^ and phosphate coexist (Metz et al. 2023). This process is most efficient in deep-sea fans and continental slopes with rapid burial of reactive FeOx and stable diagenetic conditions over thousands of years, making sub-SMT Fe(II) phosphate formation a globally relevant, yet previously underestimated, sink in the marine phosphorus cycle (März et al., 2018).

### 1.4 Methanogens and phosphate mineralization

The formation of phosphates below the SMT zone is commonly regarded as closely associated with anaerobic methane oxidation because they provide the localized Fe^2+^ enrichment, alkalinity increase, and phosphate release exactly where vivianite nucleates. Methanogens can contribute under certain conditions, but their typical metabolic footprint does not create the same hotspots for Fe^2+^-phosphate precipitation (Egger et al., 2015; Hsu et al., 2014; Slomp et al., 2013). Here we demonstrate that under specific conditions, hydrogenotrophic methanogens can form hydroxyapatite (HAp) rather than vivianite, representing a novel pathway for phosphate biomineralization (Zhuang et al., 2019). *Methanobacterium* AMB_03, isolated from the chronic phosphate-poor serpentinizing environment of Chimaera (Çıralı/Antalya/Turkey) (Neubeck et al., 2017), represents an ideal model system for this investigation: its natural low-P background provides evolutionary context for phosphate storage strategies, while controlled laboratory conditions allow mechanistic dissection of the biomineralization pathway. We show that under elevated phosphate concentrations in the presence of Ca^2+^, *M.* AMB_03 is capable of forming HAp precursors, suggesting that methanogen-driven biomineralization may provide an additional pathway for phosphate mineral formation within and below the SMT. Genomic analysis also points to the complete *ppk1*-*ppx* polyphosphate cycle as a likely genetic basis for HAp biomineralization in this strain.

## 2. Materials and Methods

### 2.1 Experimental setup

To study the growth patterns of *M.* AMB_03 under varying phosphate concentrations, two batches of continuous growth experiments were set up under anaerobic conditions, for which a liquid culturing medium was prepared across a range of phosphate concentrations (P) as shown in Fig. 1. Medium with the following six concentrations of phosphate were prepared: P = 0 mM, P = 0.1 mM, P = 0.5 mM, P = 1 mM, P = 3 mM, and P = 5.8 mM (≈ 6 mM), see Section 2.2. Blank (i.e. not inoculated) experiments were prepared for each condition. The first batch comprised four experiments per phosphorus treatment, including one “sacrifice” bottle for preliminary analysis. Based on this initial analysis, which showed high growth, a second batch was prepared with duplicate experiments per treatment, monitoring growth under conditions of energy starvation (feeding with 20%/80% CO_2_/H_2_ gas only once at the experiment setup, see 2.3). The two experimental cohorts were monitored over temporal scales of 22 and 15 weeks, respectively. To establish a comparative baseline for phosphate precipitation, control experiments were conducted using the closest known relative of *M.* AMB_03, *Methanobacterium oryzae* (DSM11106), originally isolated from a ricefield soil in Luzon, Philippines (Joulian et al., 2000), as well as the more distantly related *Methanoculleus bourgensis* MAB1 (Manzoor et al. 2016), which has been reported from both subsurface and surface settings, including biogas digesters.

**Figure 1.**
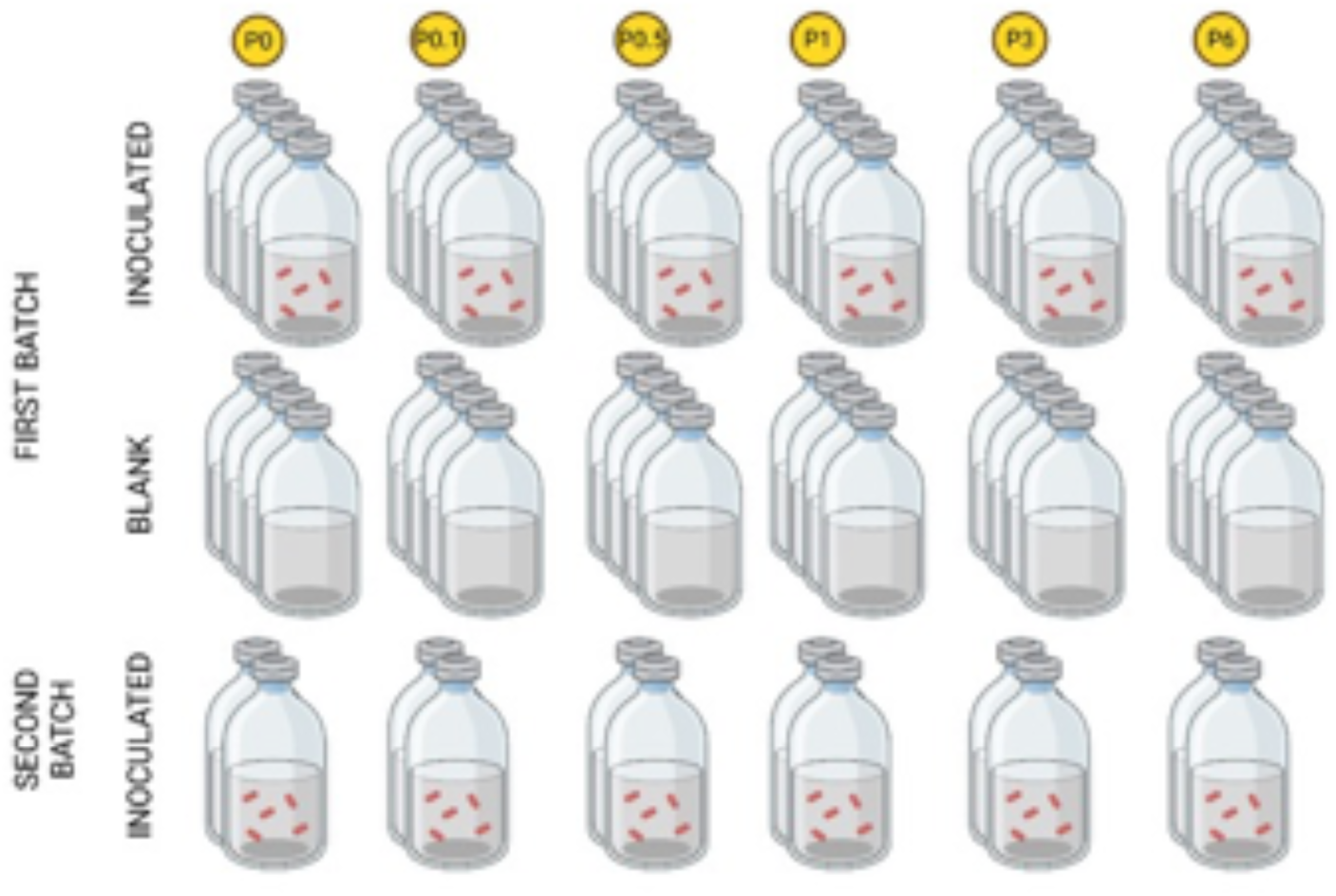
Experimental setup for two batches for cultivation of M. AMB_03 across six phosphate concentrations (P0, P0.1, P0.5, P1, P3, and P6 mM), with inoculated and blank bottles. Created in BioRender, https://BioRender.com/t2tqiex.

Both strains were grown at a phosphate concentration of 6 mM, following the identical experimental protocol described above (Sections 2.2-2.3) and initiated at a different start date to the *M.* AMB_03 experiments. *M.* oryzae was analyzed without feeding at two time points, ∼4 months (∼120 days) and 900 days, and *M.* bourgensis MAB1 at 800 days (see also Neubeck et al. 2025), prior to HAp and EPS analyses. All methanogens used for this study were pre-grown in a growth medium representing P6, see Section 2.2 below.

### 2.2 Basal medium preparation

The methanogens were grown in a bicarbonate-buffered basal medium (BM) prepared as described by Zehnder et al. (1980) and modified by Westerholm et al. (2010). Because phosphate was a controlled variable in the experimental setup, the BM was modified to be phosphate-free by excluding KH_2_PO_4_ and Na_2_HPO_4_ (solutions A and B), which were later added in different concentrations. Resazurin (final concentration 2.13 µM), Se/Wo solution (0.1 µM), and yeast extract powder (0.2 g/L; Oxoid™, UK) were measured and mixed in a large Erlenmeyer flask. Distilled water (894 mL) was added, and the liquid level was marked on the bottle. Additional distilled water was added to approximately 2 cm above the marked line. The BM was then boiled until the water level returned to the mark, allowed to cool to room temperature under continuous nitrogen gas flushing, and subsequently cooled further in an ice bath. The experimental control using *M. oryzae* and *M. bourgensis* MAB1, used an identical experiment setup as the P = 5.8 mM (≈ 6 mM) described in section 2.1 (see also Neubeck et al. 2025).

### 2.3 Sample preparation

For the first batch of experiments, forty-eight 118 mL bottles were flushed with N₂, and 18 mL of the BM (Westerholm et al., 2012) was added using an autodispenser. The bottles were then sealed with a rubber stopper and metal crimp cap, followed by three consecutive flushes with 20/80% N_2_/CO_2_, using a vacuum pump between each flush to ensure anoxic conditions. The bottles were autoclaved for 20 minutes at 121°C, cooled to 70 °C within the autoclave, and then allowed to reach room temperature.

Subsequently, 1 mL of mixture C1 (trace metals, vitamins, and salts) and 1 mL of mixture C2 (NaHCO_3_, cysteine-HCl, and Na_2_S·9H_2_O) were sterile-filtered into each bottle (Westerholm et al., 2012). Phosphate solutions were then prepared and sterile-filtered into the bottles: concentrations ranging from 0.1 to 3 mM were made using KH_2_PO_4_ (EMSURE® ISO, Germany), with 1 mL added to each bottle. For the 6 mM solution, 1 mL each of 3 mM KH_2_PO_4_ and 3 mM Na_2_HPO_4_ (EMSURE® ISO, Germany) was added. The experiments were inoculated according to Table S1.

For the second batch, consisting of twelve bottles, the same procedure was followed with one key modification: phosphate solutions were added prior to autoclaving and flushing with N_2_/CO_2_. Therefore, only the C1 and C2 mixtures were added before inoculation, as outlined in Table S1. All samples were incubated in a warm room at 37 °C and supplied with 20/80% CO_2_/H_2_ gas. Prior to feeding, gas samples were collected, and the overhead gas in each bottle was removed by piercing the rubber stopper with a needle. The first batch was fed weekly during the first five weeks and subsequently once per month, based on preliminary analysis of the sacrifice bottles. The second batch was fed only once, on the day of setup. A subset of bottles from each treatment was retained under continued incubation (37°C, 20/80% CO_2_/H_2_ feeding as described above) beyond the initial 15-and 22-week monitoring periods, and destructively sampled at week 45 (low-phosphate treatments) and week 49 (full treatment set) for SEM/EDS analysis, to capture longer-term precipitation dynamics comparable to the extended incubation periods used for the M. oryzae and MAB1 controls.

### 2.4 Genome Annotation

Genomic DNA was extracted from *M.* AMB_03 pure cultures and was sequenced by Novogene Co. Ltd (Cambridge, UK) on Oxford Nanopore R9 chemistry. Basecalling and de novo assembly were performed, followed by polishing with two rounds of Racon (Vaser et al., 2017) and Medaka (Oxford Nanopore Technologies), yielding a single closed circular contig of 2,819,821 bp. The genome was annotated with Bakta v1.11.4 (Schwengers et al., 2021) and independently with Prokka v1.14.6 (Seemann, 2014). Protein-coding genes were assigned orthology-based functional annotation (COG categories, EC numbers, KEGG orthologs) using eggNOG-mapper v2.1.9 (Cantalapiedra et al., 2021) against the eggNOG 5.0 database (Huerta-Cepas et al., 2019), with the Bakta-predicted proteins as input; these assignments are reported in Table 1.

**Table 1.** Comparative genomic analysis showing functional annotation by eggNOG-mapper (Section 2.4) experimental results.

| **Gene** | **Function** | **AMB_03** | **M. oryzae** | **MAB1** |
|---|---|---|---|---|
| **PHOSPHATE TRANSPORT** |  |  |  |  |
| *pstB, pstA, pstC* | High-affinity phosphate ABC transporter (PstSCAB) | **Present** | **Present** | **Present** |
| *PhoU-domain proteins* | Phosphate uptake regulation (archaeal-type) | **Present** | **Present** | **Present** |
| **POLYPHOSPHATE METABOLISM** |  |  |  |  |
| *# ppk1 (MCAMB03_00139–00140)* | Polyphosphate synthesis | **Present** | **Absent** | **Absent** |
| *ppk2* | Alternative polyphosphate synthesis | **Absent** | **Absent** | **Absent** |
| *ppx / Ppx-GppA (MCAMB03_00141)* | Polyphosphate degradation, canonical (PF02541) | **Present** | **Absent** | **Absent** |
| *surE* | Functional phosphatase analog (non-canonical)* | **Present** | **Present** | **Present** |
| **PHOSPHATE REGULATION & SCAVENGING** |  |  |  |  |
| *ppa* | Inorganic pyrophosphatase | **Present** | **Present** | **Present†** |
| **EPS & AMYLOID-LIKE PROTEINS (AMB_03 only)** |  |  |  |  |
| *Pseudomurein-binding repeat (M_AMB03_00342)‡* | Archaeal cell wall binding; candidate extracellular matrix scaffold | **Present** | **Absent** | **Absent** |
| *Glycosyltransferase GT39 (M_AMB03_02928)‡* | Polysaccharide synthesis; possible role in EPS production | **Present** | **Absent** | **Absent** |

*Gene presence/absence based on EggNOG annotation and BlastX searches. Green = present (canonical); red = absent; blue = present as functional analog only.*
*# Annotated as two adjacent partial ORFs that together encode a single full-length PPK1 (~713 aa); see Section 3.6.2*
*\* surE encodes a SurE-type 5'-nucleotidase (EC 3.1.3.5), distinct from canonical ppx (PF02541); exopolyphosphatase activity has not been experimentally confirmed in M. oryzae or MAB1.*
*† MAB1 encodes ppaC (DHH-type, COG1227), a structurally distinct family from the standard inorganic pyrophosphatase (COG0221) in AMB\_03 and M. oryzae.*
*‡ PLAAC-predicted candidates with COREscore > 0; see Table S8 for full list.*

### 2.5 Phylogenomic tree construction and polyphosphate gene annotation

Phylogenomic trees were constructed using GToTree v1.8.16 (Lee, 2019) with the prepackaged Archaea single-copy gene set (76 target genes). Representative genomes spanning the phylum Methanobacteriota (n = 330) were obtained from the Genome Taxonomy Database (GTDB) release r232 using the gtt-get-accessions-from-GTDB helper program with the -- GTDB-representatives-only flag. This retrieves a single representative genome per GTDB species cluster. Because GTDB defines species clusters by whole-genome average nucleotide identity (ANI) at a ∼95% threshold and designates one representative per cluster, this criterion returns every recognized Methanobacteriota species exactly once (n = 330 in r232), providing complete species-level sampling of the phylum while, by construction, excluding multiple near-identical conspecific genomes that would otherwise over-weight densely sequenced lineages and bias gene-frequency estimates. Genome selection was therefore determined by GTDB’s ANI-based dereplication rather than by manual curation. After quality filtering (genomes recovering <50% of the 76 target single-copy genes were excluded), 319 GTDB genomes were retained; *M*. AMB_03 was added as the focal strain, yielding 320 genomes for phylogenetic analysis.

Within GToTree, single-copy genes were identified using HMMER3 v3.2.2 (Eddy, 2011), individually aligned with MUSCLE v3.8.1551 (Edgar, 2004), trimmed with trimAl (Capella-Gutiérrez et al., 2009), and concatenated before phylogenetic estimation with FastTree2 v2.1.11 (Price et al., 2010). To assess the distribution of polyphosphate metabolism genes across the phylogeny, GToTree’s additional PFam search functionality (-p flag) was used to identify *ppk1* (PF02503), polyphosphate kinase 2 (*ppk2*, PF03976), and exopolyphosphatase (*ppx*, PF02541) in all genomes. Gene presence was determined using HMMER3 with an e-value threshold of 1e-5 for PFam domain hits. GToTree automatically generated presence/absence data and produced Interactive Tree of Life (iTOL) (Letunic and Bork, 2007) compatible annotation files. To supplement the HMM-based phylogenomic screen, targeted BLAST searches for *ppx* were carried out using BlastX (nucleotide query translated against a protein database), with query sequences downloaded from NCBI using the EC number 3.6.1.11 as a unifying identifier. This EC-number-based approach was adopted because *ppx* and its functional homologs carry more than 15 synonymous annotations in public databases, making name-based searches unreliable. The *E. coli* exopolyphosphatase EDU80702.1 (annotated with EC 3.6.1.11 and PF02541) was used as one reference sequence, consistent with the reference used by Paula et al. (2019). To improve the readability of the 320-genome dataset, terminal leaves were collapsed to genus level using ete3 (Huerta-Cepas, Serra and Bork, 2016), retaining one representative per genus while preserving branch lengths, resulting in 30 genus-level entries. Gene annotations at each collapsed node are displayed as proportional filled bars indicating the fraction of genomes within that genus encoding each gene.

#### 2.5.1 Targeted *ppk1*–*ppx* sequence and synteny comparison

To evaluate whether the *ppk1*–*ppx* arrangement in *M*. AMB_03 is conserved, the four complete-pathway Methanobacteriota genomes closest to *M*. AMB_03 in the phylum-wide tree (Section 2.5) were selected: two Methanobacterium_D representatives (GCA_963787255.1, and GCA_039666455.1), and two cultured *Methanosphaera* references, *M. stadtmanae* GCA_000012545.1 and *M. massiliense* GCA_028890305.1. For each genome, the *ppk1* (PF02503) and *ppx* (PF02541) protein sequences were taken from the GToTree PFam hit output, and pairwise amino-acid identity to the corresponding AMB_03 protein was calculated by global pairwise alignment with the BLOSUM62 substitution matrix. To determine gene arrangement, each assembly was retrieved from GenBank and protein-coding genes were re-predicted with Pyrodigal in metagenomic mode (Larralde, 2022), a reimplementation of Prodigal (Hyatt et al., 2010); *ppk1* and *ppx* were then located and required to reside on the same contig, and their strand, order, and intergenic distance were recorded.

### 2.6 Prediction of amyloid aggregation

We have previously demonstrated that the hidden Markov model (HMM) based algorithm, prion-like amino acid composition, PLAAC (Lancaster et al., 2014), can successfully identify amyloid-forming proteins (Zajkowski et al., 2023, 2021). The command-line version of PLAAC installed from https://github.com/whitehead/plaac was applied to all 2,945 predicted proteins in the *M.* AMB_03 proteome. We used default settings other than “-a 0”, and those with a COREscore > 0 (excluding “NA”) were considered to contain a prion-like domain. This COREscore is the sum of the individual log-likelihood ratios for each amino acid residue contained within the prion-like domain. A positive “>0” COREscore value from PLAAC is indicative of a protein’s high compositional similarity to known yeast prions.

### 2.7 Analytical methods

#### 2.7.1 Gas analyses

For the first batch, gas and pressure measurements were initially taken weekly, and every second week following the switch to monthly feeding. On a feeding day, samples were taken both before and after feeding. For the second batch, to avoid depleting the gas content in the flasks, samples were only taken for the first three weeks. Pressure measurements were taken with a GSMD 2 BR - K31 pressure sensor by Greisinger. To obtain the gas samples, 1 mL of gas was taken from the bottles using a 3 mL syringe. As the gas within the flasks is under high pressure, the syringe was removed part way to allow the gas to expand while the needle was inside the rubber stopper. After making note of the pressure and new volume, 1 mL of the uncompressed gas was inserted into the gas sample bottles. These samples were stored at-2°C before analysis. H_2_, CO_2_, and CH_4_ volume percentages were measured by light gas chromatography using a Perkin Elmer Clarus 590 configured with a TurboMatrix 110 headspace autosampler, with argon gas as mobile phase. Gas components were separated using a 7’ HayeSep N (60/80 mesh) and a 9’ Molecular Sieve 13x (45/60 mesh) followed by CH_4_ detection with a flame ionization detector, and CO_2_ and H_2_ detection using a thermal conductivity detector.

#### 2.7.2. Alcian Blue staining and imaging

Alcian blue (AB) staining was performed to test for extracellular polymeric substance (EPS) using Alcian blue solution, pH 2.5 (Sigma-Aldrich, Germany). The protocol supplied with the product was loosely followed but modified to suit the different starting material, and no counterstain was used. 1 mL of each sample was first vacuum-filtered onto a 0.20µm nylon mesh filter and rinsed once with distilled water. The filters were transferred to petri dishes, where they were submerged in the AB solution for five minutes. After staining, the filters were carefully transferred back to the vacuum filter and rinsed twice with distilled water to remove leftover AB solution. The samples were then successively washed in a series of increasing ethanol concentrations, repeating each step twice with 70%, 95%, and 100% ethanol, all the while using a vacuum pump. The filters were left to air dry and observed using regular optical microscopy.

#### 2.7.3. SEM imaging and spectroscopy

Samples were prepared for Scanning Electron Microscopy (SEM) following a two-part protocol: first fixing the cells in aldehydes as per Passey et al. (2007) then drying according to the procedure in Araujo et al. (2003). Cells were first isolated from the samples by suction filtering the samples onto a 0.2 µm nylon mesh filter. The fixing solution was prepared by mixing 4 mL of 1 M glutaraldehyde solution and 4 mL of 1 M premixed phosphate-buffered saline (PBS) buffer solution (Roche, Switzerland) in 32 mL distilled water. The filters were transferred to petri dishes where they were submerged in approximately 4 mL of the fixing solution for 4 hours. Following this, the samples were dried using an ethanol series to prevent cell lysis. The filters were removed from the glutaraldehyde + PBS solution, and washed three times in 0.1 M PBS for 10 minutes each time. They were subsequently washed using a series of increasing ethanol concentrations with 50%, 70%, 80%, 90%, and 95% ethanol for 10 minutes each, and then three times 100% ethanol for 10 minutes each. Finally, they were washed once in hexamethyldisilazane (HMDS; grade ≥ 99%; Sigma-Aldrich, Germany) for 30 seconds, after which the HMDS was removed and the filters left to air dry overnight in a fume hood.

The filters were observed under an optical microscope and areas were selected for observation. These were cut out and mounted on SEM plates using carbon conductive tabs, and were then sputter-coated with a mixture of gold and palladium. The samples were observed under SEM maintaining a working distance between 7.5 and 8.5 mm. Images were obtained at 6 kV. In addition, the chemical composition of points of interest was determined by performing a spot-analysis using energy-dispersive X-ray spectroscopy (EDS) at 20 kV. Field sediment samples from the serpentinizing site of isolation (Chimaera, Turkey, Neubeck et al., 2017) were analyzed by SEM-EDS using the same instrument settings described in Section 2.7.3. EDS mapping and spot analyses were performed to determine the elemental composition and mineral phases present in the natural sediment.

#### 2.7.4. Raman Spectroscopy and XRD analyses

Raman spectroscopy of the HAp precipitates was performed using a Horiba Scientific LabRAM HR 800 laser Raman confocal spectrometer equipped with a multichannel air-cooled charge-coupled-device (CCD) detector. An Ar-ion laser (⌊=514 nm) was used as the excitation source. The instrument was integrated with an Olympus microscope, and the laser beam was focused to a spot of 1 µm with a 100x objective and a spectral resolution of approximately 0.3 cm^-1^. The instrument was calibrated using a neon lamp and the Raman line (520.7 cm^-1^) of a silicon wafer. Instrument control and data acquisition were performed with LabSpec, version 5, software and spectra were matched against known databases such as RRUFF.

Raman analyses targeting the polyP-diagnostic (PO_2_^-^) band were performed on *M.* AMB_03 and on its closest cultured relative, *M. oryzae*, under identical acquisition settings. *M. bourgensis* MAB1 was excluded from this comparison: despite repeated attempts, filament clusters recovered from the MAB1 filters were too sparse and too small to yield a spectrum with usable signal-to-noise (Fig. S6). Because *M. bourgensis* MAB1 also lacks *ppk1* and *ppx* (Table 1) and, like *M. oryzae*, failed to precipitate HAp under identical conditions (Section 3.3), a null Raman result from MAB1 would not have added resolving power beyond what is already established by the *M. oryzae* comparison; we therefore restrict the filament-level polyP comparison to the two most closely related strains in this study, AMB_03 and *M. oryzae*, for which a genotype contrast exists on an experimental and phylogenetic background.

PolyP analyses of the methanogenic cells were carried out using a Renishaw inVia Qontor confocal Raman microscope (Renishaw plc, Wotton-under-Edge, UK) equipped with a 785 nm diode laser, a 1200 l/mm grating, and a Renishaw Centrus detector, operated via WiRE 5.6. Prior to the analysis, the instrument was calibrated against a silicon wafer (520.5 cm^-1^). A 50× long-working-distance objective was used. The spectra were acquired with a total acquisition time of 40 s for the nylon spectrum and 4 minutes for the filament spectrum, both at a laser power of ∼45 mW at the sample. The methanogen filament colonies were handpicked (after the drying method described above in Section 2.7.3) from the nylon membrane filters and placed on sterilized (99.5% ethanol) aluminum foil prior to Raman analysis. Care was taken to only select methanogen cells without visible HAp precipitates.

Phase identification was carried out on powdered samples using a Bruker D8 ADVANCE powder diffractometer with a Cu anode (0.15406 nm) and a Lynxeye XE-T PSD detector, operating at 40 kV and 40 mA, with Bragg−Brentano geometry over a 2θ range of 10 to 80° and a step size of 0.017° 2θ. Data were collected and interpreted using the software CrystalSleuth to identify the d-spacing of the main peaks, using RRUFF database and American Mineralogist Crystal Structure database (AMCSD). Data treatment (baseline subtraction and Savitzky–Golay smoothing) was performed using OriginPro 2024 (10.1.0.178).

#### 2.7.5. Growth medium analyses

Growth medium was recovered from the cell filtration and stored in Falcon tubes. Ni, Fe, Zn, Cu, P, Mo, B, Mn, Co, and Al concentrations were measured using Inductively Coupled Plasma - Optical Emission Spectrometry (ICP-OES). 3 mL of each sample were diluted with 2 mL of 5% HNO3, except for the P = 0 inoculated sample, of which 1 mL was diluted in 4 mL 5% HNO_3_ due to the previous dilution not being sufficient to yield measurements. The samples were filtered using 0.45 µm syringe filters. Avio 200 Scott/Cross-Flow Configuration was used for ICP measurements, and calibration curves were made using a multielement standard (CPAchem). A 6-point linear regression was set up for each element using concentrations of 0, 0.001, 0.01, 0.1, 1, and 5 mg/l. The growth medium contains background dissolved phosphorus from yeast extract (0.2 g/L), which may contribute to total dissolved P measurements particularly in nominally phosphate-free conditions (P0); all ICP-OES P values therefore represent total dissolved phosphorus including medium-derived background.

Additionally, pH measurements were performed using a Mettler Toledo pH meter. One experiment was selected per condition (P/inoculation), and 1 mL of the liquid was extracted, replacing the syringe each time to avoid cross-contamination. Iron was added to the growth medium as FeCl_2_*4H_2_O (MW 198.81 g/mol; ferrous chloride tetrahydrate), giving a nominal Fe(II) concentration of 10.06 µM. Under the anaerobic, sulfide-containing medium conditions, partial precipitation of Fe as iron sulfide (mackinawite, FeS) is expected prior to inoculation, and the measured dissolved Fe concentrations (ICP-OES, ∼1.4–2.2 µM across P treatments) reflect the fraction remaining in solution after this initial equilibration, rather than the nominal added amount.

#### 2.7.6 Thermodynamic modelling

Saturation indices (SI) for HAp, calcite and vivianite were calculated using PHREEQC version 3 (Parkhurst and Appelo, 2013) with the phreeqc.dat thermodynamic database. Input solution chemistry for each solution was defined using measured endpoint values: Dissolved phosphorus concentrations from ICP-OES endpoint measurements were converted from mg/L to mM by dividing by the atomic mass of phosphorus (30.974 g/mol). Calcium was fixed at 0.75 mM (7.5×10^-4^ M) corresponding to the CaCl_2_*2H_2_O content of the basal medium (Zehnder et al., 1980; Westerholm et al., 2010). Dissolved Ca was not measured by ICP-OES and was therefore held at the nominal medium concentration (0.75 mM) for all PHREEQC calculations. Biological Ca uptake into precipitates in high-precipitation conditions may have lowered true endpoint dissolved Ca relative to this nominal value, meaning the modelled HAp saturation indices in heavily-mineralizing bottles should be treated as upper-bound estimates. Dissolved Fe from ICP-OES endpoint measurements was treated as Fe(II) under the anaerobic experimental conditions and used for vivianite saturation calculations. The nominal Fe(II) added to the medium (10.06 µM as FeCl_2_*4H_2_O) substantially exceeds the measured dissolved Fe (∼1.4–2.2 µM), consistent with partial FeS precipitation prior to sampling; the measured values were therefore used as model inputs to reflect actual solution chemistry at the time of analysis. Bicarbonate alkalinity was set to 47.6 mM, consistent with the NaHCO_3_ content of the growth medium. Temperature was set to 37°C, matching incubation conditions. Saturation index is defined as SI = log(IAP/Ksp), where IAP is the ion activity product and Ksp is the solubility product; SI > 0 indicates supersaturation. Bicarbonate alkalinity was fixed at the nominal medium value of 47.6 mM; while methanogenesis consumes dissolved CO₂ and the actual carbonate system will have evolved over 22 weeks, measured pH increases in inoculated bottles (Table S4) confirm that the net effect is alkalinity increase rather than decrease, supporting the model’s qualitative conclusions.

## 3. Results

### 3.1 Gas data

There is an increase in CH_4_ with time in both the first and second batch experiments (Fig. 2).

**Figure 2.**
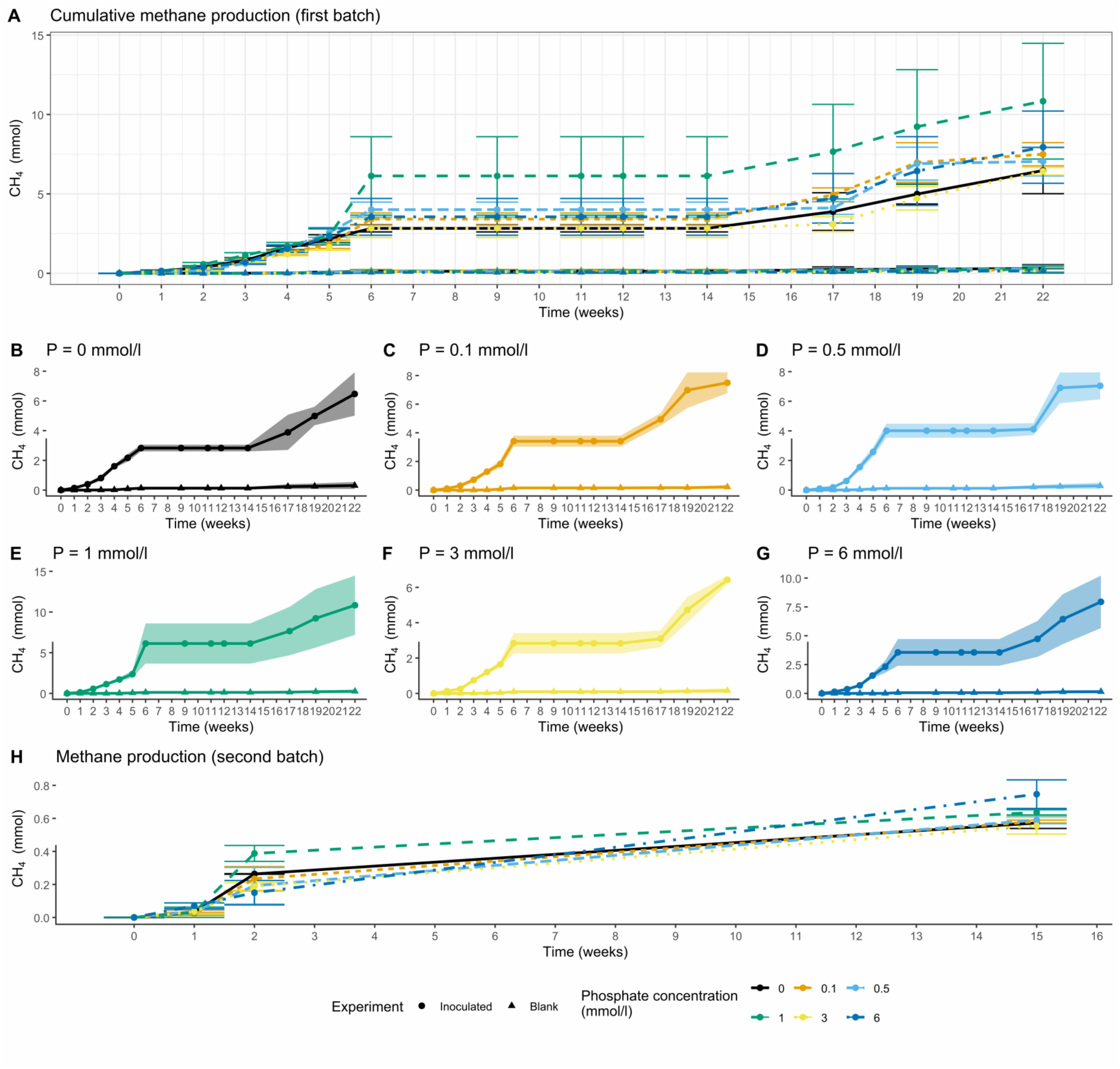
A) Cumulative CH_4_ production (mmol) across all phosphate treatments in the first batch of experiments (overlay of B-G). B) P=0 mM. C) P=0.1 mM. D) P=0.5 mM. E) P=1 mM. F) P= 3 mM. G) P=6 mM. H) CH_4_ concentration (mmol) over time in the second batch (all treatments overlay). Error bars correspond to the standard error of the mean.

**Figure 3.**
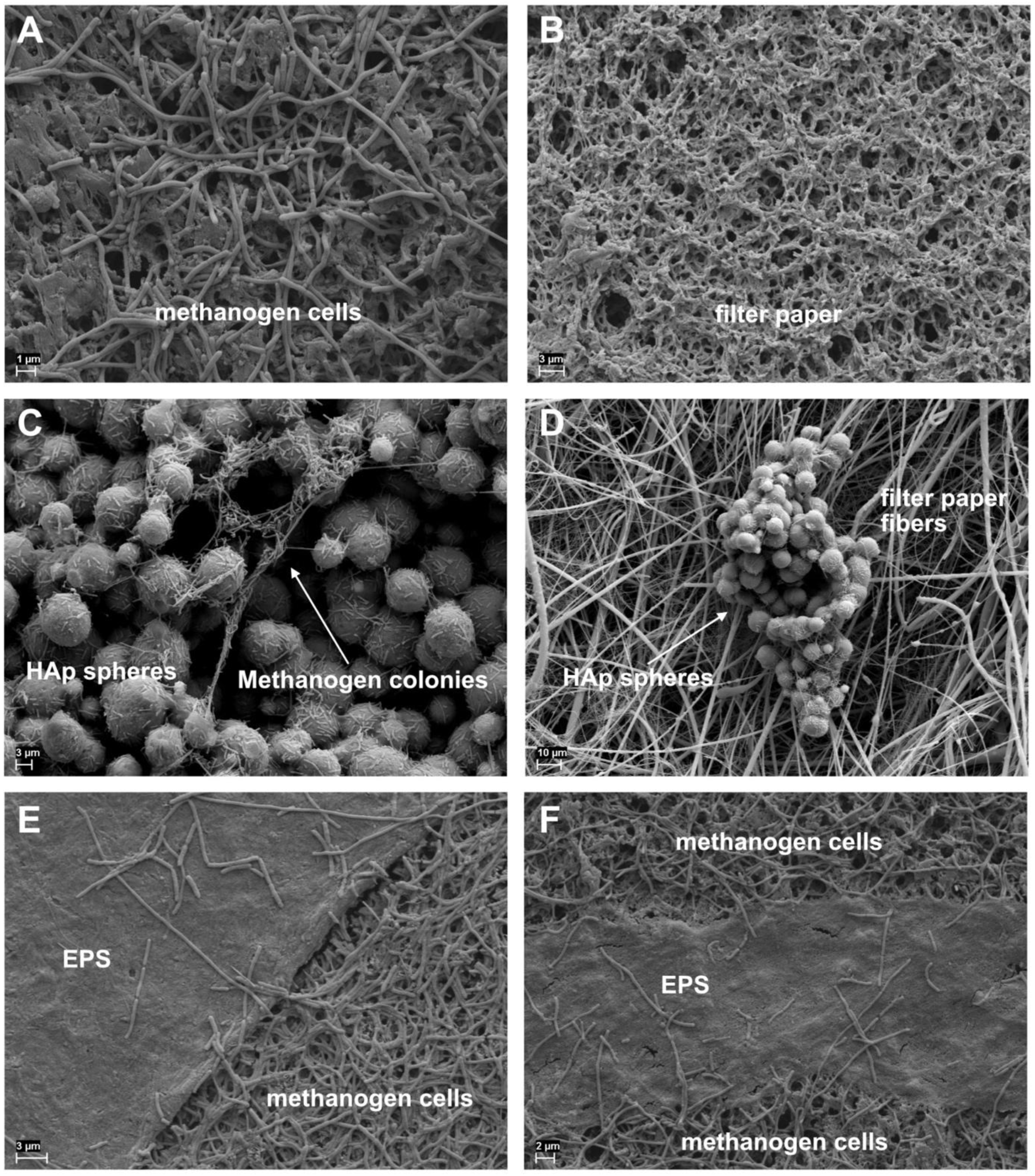
SEM images showing a) abundant cells, b) no precipitates in the blank, c-d) spherical precipitates and e-f) sheet-like EPS.

Due to leakage during gas storage, CH_4_ measurements collected between weeks 9 and 14 could not be reliably quantified. Although CH_4_ production likely continued during this interval, the affected measurements are characterized by an artificial plateau in the accumulation curve. To prevent the inclusion of potentially biased values, we conservatively assigned zero CH_4_ production to these sampling points. The time points were retained in Fig. 2 to preserve the continuity of the experimental record and to explicitly indicate the period impacted by sampling artifacts. In both batches, the lower phosphorus concentrations exhibit higher initial CH_4_ production. However, as the experiments progressed, a shift occurred in which the higher-phosphorus treatments ultimately produced more CH_4_. Methane production in the second batch was consistently an order of magnitude lower than in the first batch across P concentrations. After 15 weeks, methane ranged between 0.50 to 0.83 mmol, whereas the first batch reached 3.43-10.69 mmol after 22 weeks, and had already exceeded 1 mmol in all inoculated samples by week 4. No significant correlation was found between CH_4_ production and phosphate concentration in the first batch, but the second batch showed a significant, moderate positive correlation (Fig. S1). Pearson correlation for the first batch was r =-0.134 at the 6-week mark (p = 0.8) and r = 0.009 at 22 weeks (p = 0.972), whereas the second batch showed r = 0.617 (p = 0.033) at 15 weeks.

### 3.2 EPS observation

Alcian Blue staining was performed to identify the potential presence of EPS (Fig. S2). The microscope magnification was insufficient to resolve individual cells, but clear differences were observed between the blank and inoculated samples. The blank showed faint, patchy staining with largely unstained background and in contrast, the inoculated samples displayed more numerous and intense dark-blue spots, with occasional filamentous structures.

### 3.3 SEM and EDS

A very high concentration of cells across inoculated experiments was observed (Fig.3a), and an absence of cells in the non-inoculated control experiments (Fig.3b). The opened bottle of the P = 0.5 experiment, however, showed signs of contamination, and was thus removed from further analyses (Fig. S3). No signs of contamination were observed in the other samples. Two types of solid phases were observed in the biotic samples, spherical precipitates (Fig.3c-d) and EPS (Fig.3e-f, See section 3.2, Fig. S4).

As established in the control experiments (section 2.1; Fig. S5; Neubeck et al. 2025), no precipitates were observed for *M. bourgensis* MAB1 even after 800 days. No precipitates were likewise observed in the *M. oryzae* after analysis at *i)* ∼4 months and *ii)* at 900 days.

The spherical precipitates occurred in the high-phosphate experiments with AMB_03 (P3 and P6) within the experiment duration time (15 weeks), but when analyzed after an additional 30 weeks (after a total of 45 weeks of incubation) spheres were also found in the low phosphate experiments (P0.1, P0.5 and P1). The diameter of the spheres varied between a few micrometers to millimeters in size, depending on the time of incubation. Many spheres showed one or more perforations with a diameter closely matching that of individual methanogen cell diameters, and in several cases the methanogen filaments were observed extending directly through these perforations rather than resting on the sphere surface (Fig.4a-b). This close dimensional correspondence between perforation size and cell diameter, combined with the consistent spatial association between perforations and methanogen cells across multiple spheres, argues against these features being artefacts of the dehydration and HMDS drying procedure (Section 2.7.3). Fixation-related shrinkage or cracking would be expected to produce irregular, randomly distributed fissures unrelated to cell dimensions, rather than discrete, circular perforations consistently sized to match methanogen cell diameter and consistently occupied by cells. Some precipitate spheres were growing directly on the cells, whereas some were covering the cells (Fig.4c-d). Calcium carbonates were observed in the low phosphate experiments, P0 and P0.1 (Fig. 4e-f).

**Figure 4.**
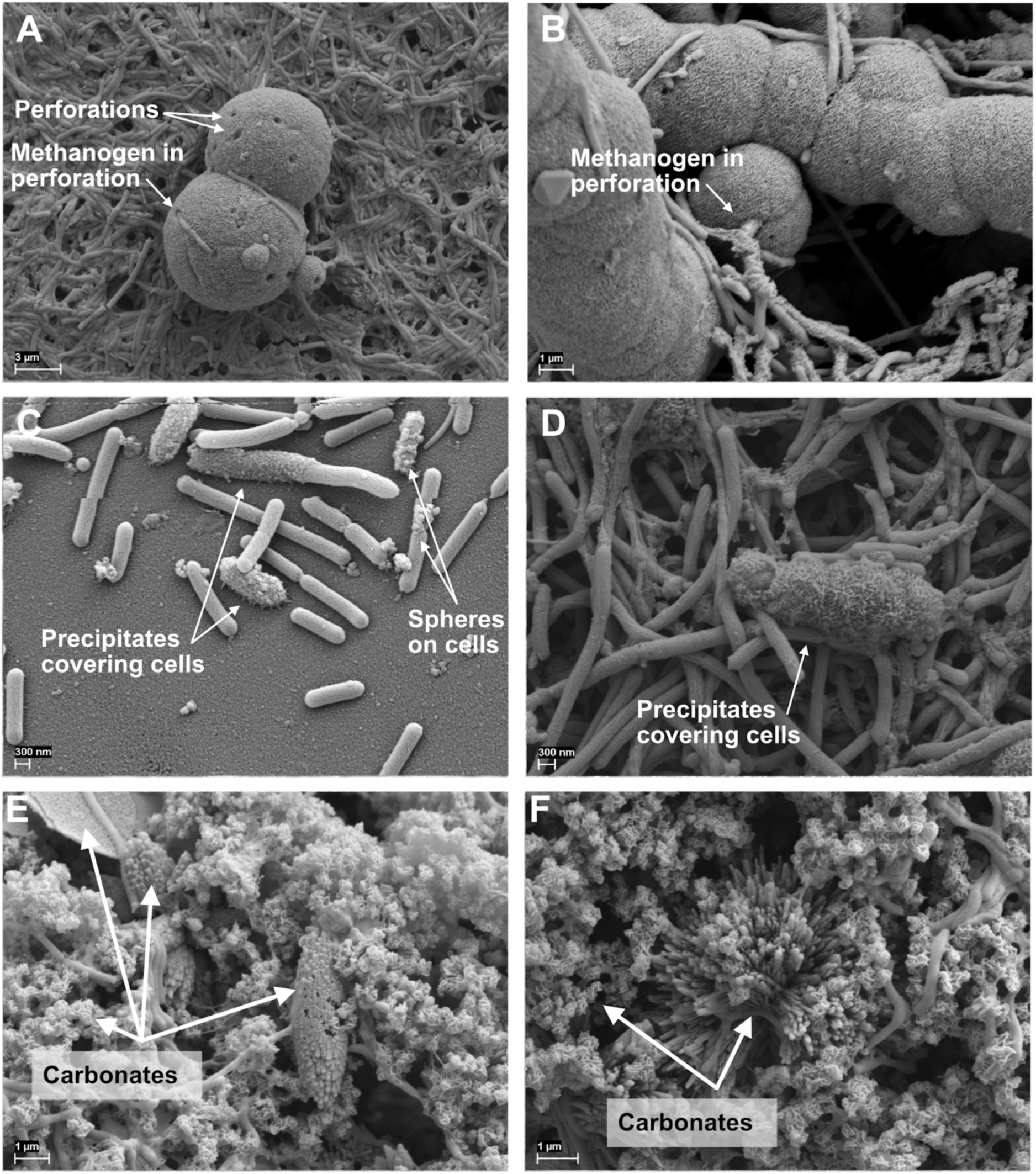
SEM images showing a-b) perforations on the spheres and methanogens growing into the perforations, c-d) precipitates forming on and around the cells, e-f) carbonates with different morphologies in the low P experiments (P0, P0.1).

No calcium phosphate precipitates were observed at lower phosphate concentrations within the first 15 weeks of the experiment duration. Likewise, none were detected at any phosphate concentration during a preliminary analysis of one sacrificed bottle per setup at week 6. Precipitation only appeared to begin after feeding with H_2_/CO_2_ was reduced to once a month, and after 49 weeks all inoculated bottles (except P0) showed calcium phosphate precipitation.

The carbonate morphologies vary from very small spheres (< 500 nm) with a rose-like texture, larger pointed ovals (up to at least ca. 10 µm in length), calcite spikes arranged in a dumbbell-like formation, as well as very large flakes. These were found growing around cells of *AMB_03* as well as amorphous organic matter. Pure carbonates were not observed in the other inoculated methanogen experiments (*M.oryzae* and MAB1), nor in the blank experiments. Like the calcium phosphates, the calcium carbonates were also not observed during the preliminary analysis at week 6. The only precipitation observed in the blank experiments were zinc sulfides (Table S2), which are chemically and morphologically very distinct from the calcium phosphate and calcium carbonate observed in both the biotic and white sediment samples. EDS analyses of the white sediment field samples from the AMB_03 isolation site revealed exclusively carbonate phases; no phosphate-bearing minerals were detected (Table S2).

### 3.4 Raman spectroscopy and XRD

The main Raman peak of the spherical precipitates (P1, P3 and P6) was located at ∼960 cm^-1^ and with three weak, broad peaks at ∼428, ∼590 and ∼690 cm^-1^ (Table S3, Fig. S6). A band at 1070–1072 cm⁻¹ was also present. The bands at ∼428 and ∼590 cm⁻¹ correspond to the ν₂ and ν₄ PO₄³⁻ modes of HAp, whereas the weak bands at ∼690 and ∼1301–1305 cm⁻¹ (Table S3) are not attributable to HAp; the latter most likely reflects organic CH₂ twisting. Raman spectra collected from EPS within the same samples showed a distinct set of peaks, with peaks at 953.5, 1064.0, 1129.0, 1235.0, 1296.5, 1442.5 and 1632.5 cm⁻¹ (Fig. S6). The average I₁₀₇₀/I₉₆₀ intensity ratio of the spheres was 0.19.

Raman spectra of methanogen filaments of M. AMB_03 selected to exclude cells with visible HAp precipitates, showed a peak at ∼1177 cm⁻¹ (Fig. 5a). To test whether this putative polyP signal tracks genotype, filaments of M. oryzae, the closest cultured relative of M. AMB_03 and, like M. bourgensis MAB1, a strain lacking *ppk1* and *ppx* (Table 1), were analyzed. Across two replicate M. oryzae filament spectra, no peak was resolved in the 1145–1177 cm⁻¹ polyP window (Fig. 5b). *M. bourgensis* MAB1 filaments could not be analysed by Raman: cell clusters recovered from the filters were too few and too small to generate a spectrum above the noise floor (Fig. S6).

**Fig. 5.**
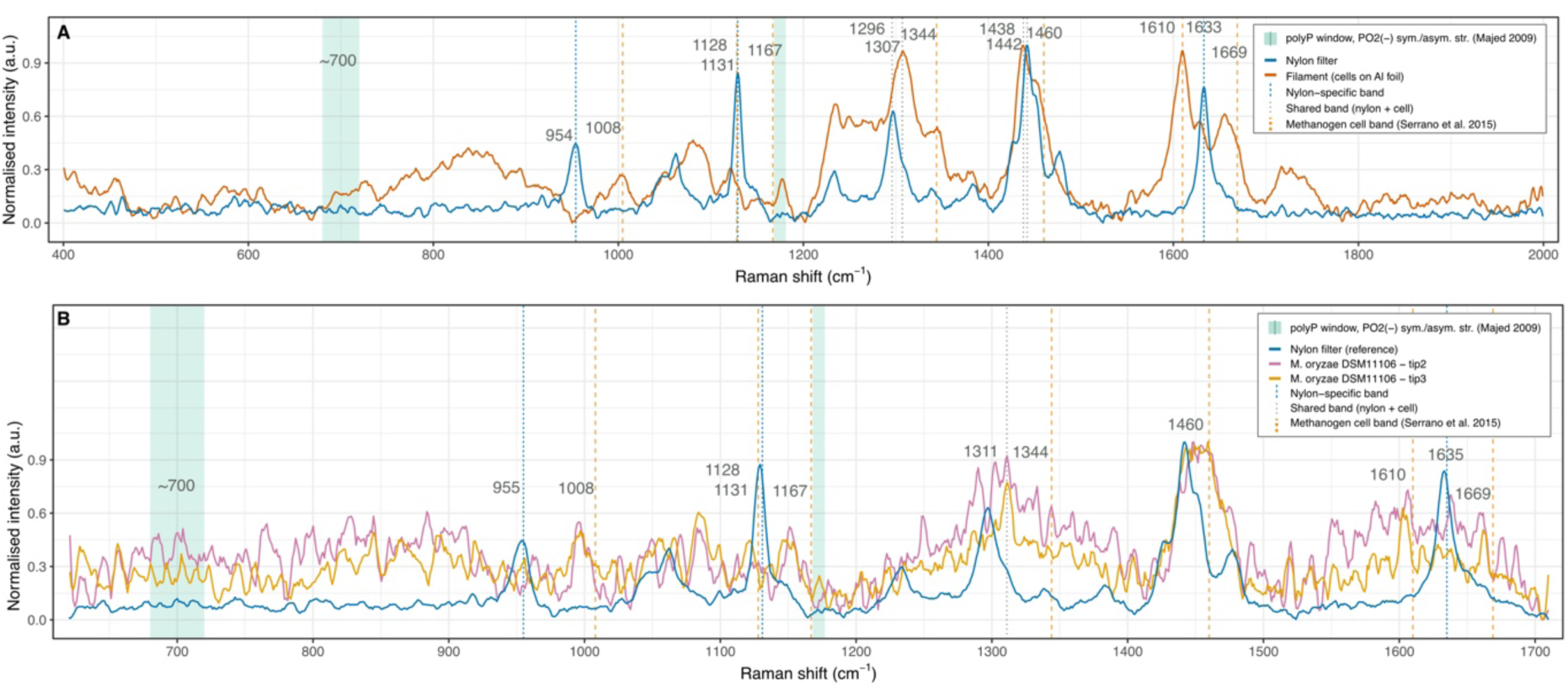
Raman spectra of M. AMB_03 filaments and a pure nylon filter (a), and of M. oryzae filaments (b). Data treatment included cosmic ray removal, baseline subtraction and smoothing. Reference peak identifications are found for polyP in Majed et al. 2009, for nylon filters in Miller & Bartick, 2001 (after Hendra et al. 1990), for methanogenic cells in Serrano et al. 2015 and other organic bands in Czamara, K. et al. (2015) and Pezzotti, G. (2021).

X-ray diffraction of the Ca-P precipitates produced a diffractogram with peaks corresponding to ten of the twelve HAp peaks matched against the Joint Committee of Powder Diffraction Standards, (JCPDS) 09-0432 reference pattern (Fig.6). Only the two most intense reflections (∼26° and ∼32° 2θ) are clearly resolved above baseline; the remaining eight peaks are broad, low-amplitude features consistent with poorly crystalline material. The two most intense peaks occurred at approximately 26° and 32° 2θ, corresponding to the (002) and overlapping (211)/(112)/(300) peaks, respectively. Additional peaks were detected at approximately 29°, 34°, 39°, 49° and 53°, corresponding to the (102)/(210), (202), (310), (213), and (004) planes. Because the calcite (104) reflection at 29.4° 2θ coincides with the HAp (102)/(210) reflections, a minor calcite contribution to the ∼29° peak cannot be excluded. Peak widths were broad throughout the pattern (Fig.6). XRD confirmed calcite as the dominant carbonate phase (Fig.S7), with the diagnostic (104) reflection at 29.4° 2θ and secondary reflections at 23.2°, 36.2°, 39.6°, 43.4° and 61.0° 2θ. Minor peaks at 27.3° and 31.7° 2θ are consistent with trace halite (likely evaporitic residue from the growth medium).

**Fig. 6.**
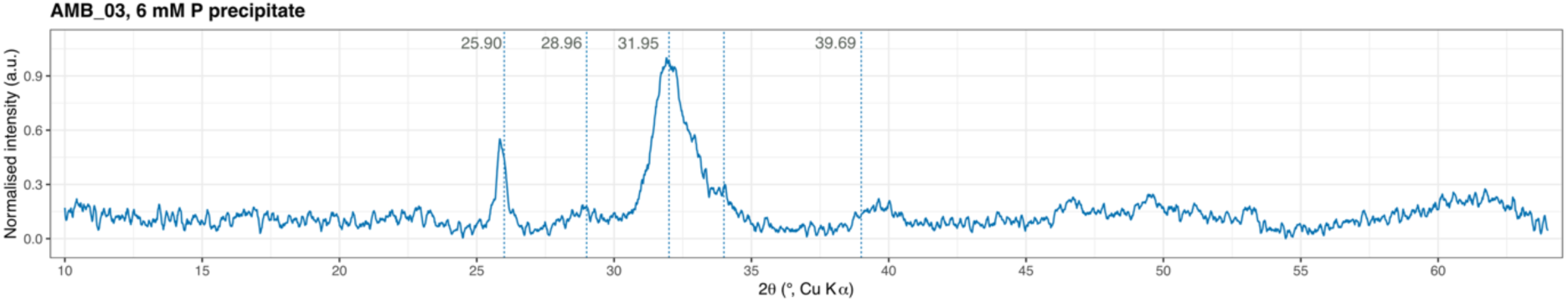
The XRD pattern of Ca-P precipitates from AMB_03 cultures. Full diffractogram (10–80° 2θ, Cu Kα radiation) with fully resolved HAp reference peak positions indicated by dashed lines plus numerical values (JCPDS 09-0432; Kay et al., 1964).

### 3.5 Growth medium analyses

#### 3.5.1 ICP-OES

The pH was measured at the start (pH ∼7) and from one sample per condition at week 25. The pH was relatively consistent across the P concentrations in the blank experiments and overall higher in all the inoculated experiments (Table S4).

The ICP-OES metal analyses were conducted on the inoculated samples, the blanks, and the growth media at the differing P concentrations (Fig. 7). Copper measurements were excluded due to their relative standard deviation (measurement spread) exceeding the recommended 6% threshold. For Ni, Mo, Mn, Co, and P, the concentrations in the inoculated samples were consistently lower than in the abiotic controls. With the exception of Fe, Zn, and P, the blanks showed minimal variation across the P concentrations, whereas both the inoculated samples and growth media frequently exhibited decreasing concentrations with increasing P. The inoculated AMB_03 samples showed little variation in Co, Ni, and Zn across the P gradient (Fig. 7). Biological phosphorus drawdown, calculated as the difference between growth medium P and endpoint dissolved P, increased with initial phosphate concentration in inoculated experiments relative to blank controls. At P0.5 and above, the non-inoculated blanks samples had higher levels of dissolved phosphorus compared to samples from inoculated bottles, with the largest biological excess removal observed at P6 (+0.25 mM relative to the abiotic control). At P0 and P0.1, blank bottles showed greater net P loss than inoculated samples, likely reflecting abiotic surface adsorption or trace precipitation, compounded by background dissolved phosphorus from yeast extract in the growth medium, which is non-negligible relative to the low nominal P concentrations in these conditions.

**Figure 7.**
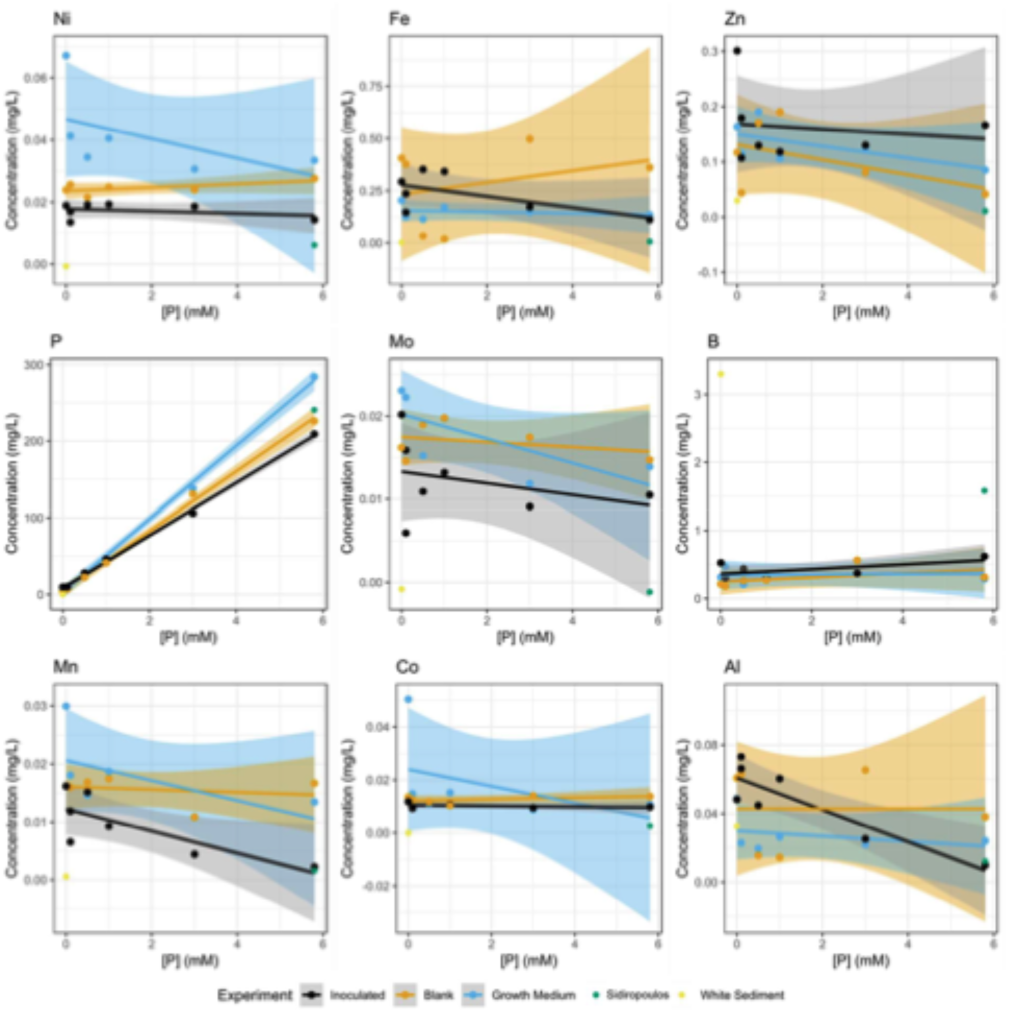
Metal concentrations in the inoculated experiments (black), blank experiments (orange), and untreated growth medium (blue) are plotted against the phosphate concentration of the prepared growth medium, with linear regressions applied. Shaded areas indicate the standard error of the regressions. Data from a white carbonate-bearing sediment sample from the AMB_03 isolation site (Neubeck et al. 2017).

Biological phosphorus decrease showed a positive directional trend with cumulative CH_4_ production across the four conditions where biological P removal exceeded abiotic loss (P0.5–P6; n = 4), though with only four data points this is reported descriptively. P0 and P0.1 were excluded from this comparison because yeast-extract background phosphorus dominates the P budget at these nominal concentrations (Section 2.7.5), not because they were outliers to the trend. Iron concentrations at the experimental endpoint showed a pattern distinct from all other measured metals. In blank bottles, dissolved Fe at several P concentrations (P0.1, P0.5, P3) was significantly higher, indicating net Fe release into solution. In contrast, dissolved Fe in inoculated bottles was consistently below growth medium concentrations across all P treatments. Mo and Ni showed the highest proportional depletion in inoculated samples relative to growth medium (∼88% and ∼64% respectively, averaged across P treatments).

#### 3.5.2 PHREEQC modelling

Saturation indices (SI) for HAp, calcite and vivianite were calculated for each phosphate treatment using PHREEQC (Table S5). Nominally phosphate-free (P0) solutions retained measurable dissolved phosphorus from yeast-extract background (see Section 2.7.5) and were included in the modelling on this basis. Inoculated solutions were supersaturated with respect to HAp across all phosphate concentrations including P0 (SI = +5.6 to +8.8), as were blank solutions (SI = +1.2 to +5.4). Inoculated solutions were also supersaturated with respect to calcite (SI = +1.1 to +1.4), as were blanks (SI = +0.3 to + 0.6). All solutions, inoculated and blanks, returned negative saturation indices for vivianite (SI =-0.8 to-3.5).

### 3.6 Genomic analyses

#### 3.6.1 Phosphate uptake and transport genes

All three strains encode phosphate ABC transporter components (*pstB*, *pstA*, *pstC*), confirmed by EggNOG annotation, indicating that differential phosphate uptake capacity does not explain the contrasting biomineralization phenotypes (Table 1). Phosphate scavenging capacity is similarly conserved: all three strains encode *ppa* (inorganic pyrophosphatase) and PhoU-domain regulatory proteins (Table 1), indicating that upstream phosphate handling does not account for the differential biomineralization phenotype.

#### 3.6.2 Polyphosphate metabolism genes

A defining genomic distinction is AMB_03’s unique polyphosphate metabolism capacity. AMB_03 lacks *ppk2* but encodes *ppk1* and *ppx*, if expressed enabling robust polyphosphate synthesis from ATP and degradation. In contrast, both *M. oryzae* and MAB1 completely lack *ppk1* and *ppk2*. Regarding *ppx*: HMM-based screening (PF02541) identified a canonical *ppx* ortholog exclusively in AMB_03, co-localized with *ppk1* in a conserved *ppk*-*ppx* gene pair (AMB03_00139-00141). In *M. oryzae* and strain MAB1, BlastX searches against an EC 3.6.1.11 protein database retrieved *SurE*-type proteins that carry this EC annotation and share functional overlap with PPX, but belong to a distinct protein family and lack the canonical *ppk*–*ppx* operon context. These proteins may represent functional analogs capable of exopolyphosphatase activity, but experimental confirmation would be required to establish this. The *ppk1* gene is annotated as two adjacent partial ORFs (MCAMB03_00139/00140) that together encode a single full-length PPK1 (∼713 aa); the split coincides with a homopolymer region and is consistent with a basecalling/assembly artifact rather than a gene duplication. It is immediately adjacent to *ppx* (MCAMB03_00141), forming a conserved *ppk1*–*ppx* gene pair.

#### 3.6.3 Phylogenetic distribution of polyphosphate metabolism genes

A phylogenomic survey of the 320 *Methanobacteriota* genomes (319 GTDB r232 species representatives plus *M.* AMB_03; Fig. 8) was screened for *ppk1*, *ppk2* and *ppx*. *ppk1* was detected in 24.1% of genomes (77/320), *ppk2* in 15.3% (49/320) and *ppx* in 33.4% (107/320; Table S6a). A complete polyphosphate cycle of at least one synthesis gene (*ppk1* and/or *ppk2*) together with *ppx* was present in 24.7% of genomes (79/320). Of the 107 *ppx*-containing genomes, 28 (26%) lacked a synthesis gene. *M.* AMB_03 encoded *ppk1* and *ppx* but not *ppk2* (Fig. 8). These genes were not randomly distributed across the phylogeny but showed pronounced lineage-level structure (Fig. 8). The three genes also partitioned into distinct clades: *ppk2* was essentially restricted to the genera *Methanocatella* (42/98 genomes) and *Methanobrevibacter* (6/25), occurring in only 3 of the 30 genera sampled (Table S6b) whereas *ppk1* and the complete *ppk1*-*ppx* cycle were concentrated in a subset of *Methanobacterium*-related genera, e.g. *Methanobacterium_A* (14/19 genomes with a complete cycle), *Methanobacterium_B* (15/30), *Methanobacterium_F* (4/4), *Methanarmilla* (7/8) and UBA349 (5/6). Conversely, the thermophilic *Methanothermobacteraceae* (0/9) and several genera (*Methanothermobacter*, *Methanovirga*) lacked all three genes. The subset of genomes carrying *ppx* without any synthesis gene was itself lineage-linked, being proportionally most concentrated in *Methanosphaera* (*ppx* in 17/23 genomes but *ppk1* in only 5). The *M.* AMB_03 itself falls within GTDB genus *Methanobacterium_D*, the same genus as the pathway-negative comparator *M.* oryzae which has the lowest complete-pathway frequency among the *Methanobacterium* splits with more than one representative genome (2/22, 9%); *M.* AMB_03’s intact *ppk1*–*ppx* cycle is therefore uncommon within its genus (present in only ∼9% of other *Methanobacterium_D* representatives) rather than typical of it. The phylogenetically clustered, clade-specific distribution of these genes indicates that polyphosphate-cycling capacity in the *Methanobacteriota* has been shaped by lineage-level gain/loss rather than being uniformly distributed.

**Figure 8.**
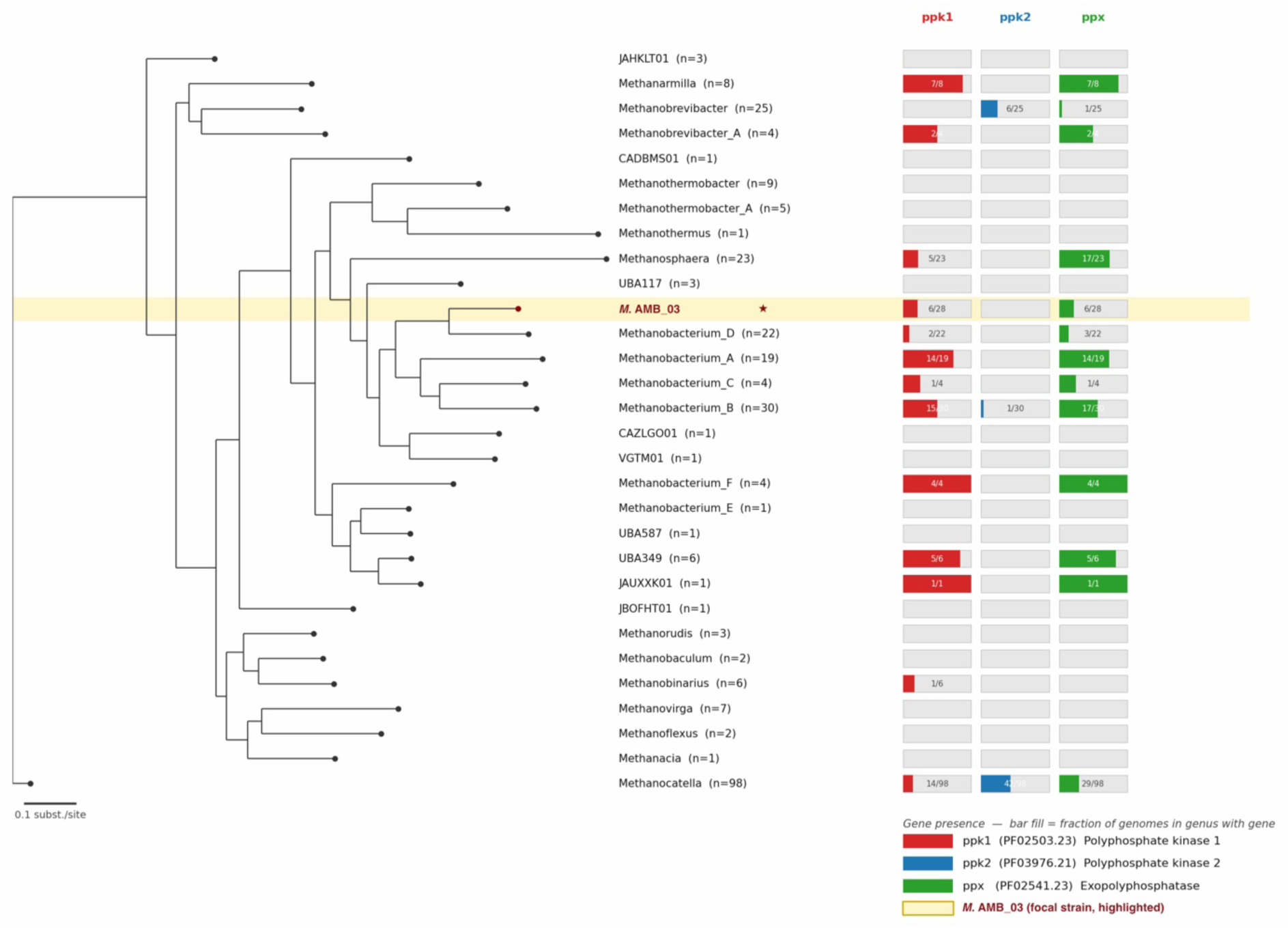
Phylogenetic distribution of polyphosphate metabolism genes across Methanobacteriota. Phylogenomic tree of 320 representative Methanobacteriota genomes (319 from GTDB release r232; M. AMB_03 included as the focal strain) constructed from 76 archaeal single-copy genes using FastTree2 v2.1.11. Terminal leaves are collapsed to genus level (30 genera); coloured bars at each genus indicate the fraction of genomes within that genus encoding polyphosphate kinase 1 (ppk1, PF02503.23, red), polyphosphate kinase 2 (ppk2, PF03976.21, blue), and exopolyphosphatase (ppx, PF02541.23, green). M. AMB_03 (highlighted, yellow) encodes both ppk1 and ppx but not ppk2, placing it among the ∼25% of Methanobacteriota genomes with complete polyphosphate cycling capability. M. oryzae and M. bourgensis MAB1 lack canonical ppk1, ppk2 and ppx orthologs; SurE-type proteins with EC 3.6.1.11 activity were detected in M. oryzae and MAB1 by BlastX but are not considered canonical ppx (see Section 3.6.2). Scale bar = 0.1 substitutions per site.

## 4. Discussion

### 4.1 Characterization of precipitates in the AMB_03 experiments

Raman spectroscopy confirms carbonate-substituted HAp (Fig. S6, Table S3). The I₁₀₇₀/I₉₆₀ intensity ratio of 0.19 indicates carbonate substitution (Gemeri et al., 2023), but because the ∼1070 cm⁻¹ region also contains a PO₄³⁻ ν₃ contribution and no calibration against standards of known carbonate content was made (cf. Awonusi et al., 2007), we do not convert this ratio into a carbonate weight fraction. The main phosphate peak at 960 cm⁻¹ lies just outside both the carbonate-substituted apatite range (955–959 cm⁻¹) and the pure crystalline HAp range (962–964 cm⁻¹) reported in the literature, falling in the narrow transition zone between them, consistent with a partially carbonate-substituted, poorly crystalline phase rather than a precise match to either end-member. The 1070/1072 cm^-1^ band confirms the CO_3_^2-^ C-O stretch. Carbonate substitution is typical of biologically precipitated apatite formed under low-temperature aqueous conditions (Mänd et al., 2018). This contrasts with methanogenic sediments, where vivianite rather than HAp has been reported to form under comparable anoxic conditions (Egger et al., 2015). We interpret this difference as evidence that the phosphate mineral phase produced by methanogens depends on both strain-specific biological capabilities and the local geochemical environment, rather than on anoxic methanogenic metabolism alone. HAp identity is further corroborated by EDS co-localization of P and Ca in the precipitates (Table S2). Ca/P atomic ratios in spheres explicitly identified as HAp precipitates (0.75-1.26; Table S2, rows labeled “HAp sphere” or “HAp+cells”) range from moderately to substantially sub-stoichiometric relative to pure HAp (1.67). The lower end of this range (0.75) is below what carbonate substitution alone typically produces; a minor amorphous calcium phosphate or octacalcium phosphate (Ca/P = 1.33) component cannot be excluded, whereas brushite (Ca/P = 1.0) is unlikely because its ν₁ band at ∼985 cm⁻¹ was not observed. The upper end (1.26) is consistent with carbonate-substituted, poorly crystalline HAp as indicated by XRD (Fig. 6). Quantitative EDS on sub-micrometre to micrometre-sized spheres resting on filters or organic material is, however, prone to systematic errors in Ca/P from sphere geometry, beam penetration into the substrate and organic-matrix effects, so the low ratios more likely reflect analytical limitations than a non-apatitic phase. Together with the Raman ν₁ band at 960 cm⁻¹ and the apatite XRD reflections, the precipitates are therefore best identified as poorly crystalline, carbonate-substituted HAp, with amorphous or octacalcium phosphate-like material at most a minor component. A second spectral type (953-1632 cm^-1^) overlaps with EPS co-occurring with the Ca-P phases, discussed in Section 4.2. Broad peak widths are consistent with poorly crystalline, nanocrystalline HAp typical of biogenically precipitated apatite (Ensikat et al. 2016).

The intracellular polyP assignment at ∼1177 cm^-1^ (Fig 5), seen in *M.* AMB_03 Raman spectra is supported by its separation from nylon’s C–C backbone stretch at ∼1123-1126 cm^-1^, which lies below the established polyP PO_2_^-^ range at 1145-1177 cm^-1^ (Majed et al. 2009). The absence of diagnostic nylon peaks from the filament spectrum rules out nylon as the source of ∼1177 cm^-1^ peak but does not exclude alternative cellular contributions. The most likely alternative is the ring C–H bending band of phenylalanine and tyrosine in proteins (∼1175–1180 cm⁻¹); however, proteins are present in both strains, whereas the band was observed only in *M.* AMB_03, which argues against a protein origin. The genotype contrast between *M. oryzae* and *M.* AMB_03 is reinforced by comparing the filaments of *M. oryzae* and *M.* AMB_03: the M. oryzae filaments showed no corresponding peak in the polyP window, consistent with the absence of intracellular polyP synthesis machinery. We did not obtain an equivalent measurement from *M. bourgensis MAB1* due to insufficient biomass on the filters; however, since *M. bourgensis MAB1* shares the same *ppk1/ppx*-negative genotype as *M. oryzae* and shows the same absence of HAp precipitation (Section 3.3), the AMB_03 - *M. oryzae* contrast already isolates the variable of interest (polyP synthesis capacity), while controlling for phylogenetic proximity, which the more distantly related *M. bourgensis MAB1* would not have improved on even had a spectrum been obtainable.

### 4.2 EPS

The EPS Raman spectral type, with the 930-970 cm^-1^ band reflecting OH···O vibrations in organic acids, the ∼1442 cm⁻¹ band CH₂ deformation, and the ∼1632 cm⁻¹ band amide I and/or H₂O bending (De Gelder et al., 2007), is independently confirmed by Alcian blue staining (Fig. S2) and SEM imaging (Fig. S4, Table S3). The 953.5 cm⁻¹ band also coincides with the ν₁ PO₄³⁻ band of amorphous calcium phosphate (∼950–955 cm⁻¹), so a contribution from an amorphous Ca-P precursor within the EPS cannot be excluded; this would be consistent with EPS acting as a nucleation site. Its spatial association with Ca-P phases implicates organic matrices in mediating nucleation (Weiner and Addadi, 1997). The production of EPS by *M.* AMB_03, together with biotically driven pH increases (7.88–8.08, Table S4), creates localised microenvironmental conditions conducive to HAp nucleation. Hydrogenotrophic methanogenesis consumes dissolved CO_2_ (CO_2_ + 4H_2_ → CH_4_ + 2H_2_O), shifting the bicarbonate equilibrium toward higher carbonate alkalinity and rising pH, explaining why pH increase scales with methanogenic activity rather than phosphate concentration. HAp precipitation is highly sensitive to pH because saturation state increases non-linearly as both hydroxide activity and phosphate deprotonation are pH-dependent (Inskeep and Silvertooth, 1988). As shown in Section 3.5.2 (Table S5), both inoculated (SI = +5.6 to +8.8) and blank (SI = +1.2 to +5.4) solutions were strongly supersaturated with respect to HAp. The high inoculated values reflect the combined effect of elevated pH and phosphate activity. The absolute SI values should be treated with some caution, as they approach the upper reliability range of HAp thermodynamic parameters in the phreeqc.dat database; even so, they remain a robust indication of the qualitative pattern of strong supersaturation when inoculated versus weaker but present supersaturation in blanks. Yet blanks showed no precipitation, consistent with kinetic inhibition in the absence of biological nucleation surfaces such as EPS (Dupraz et al., 2009). *M. bourgensis* MAB1 and *M. oryzae*, which also conduct hydrogenotrophic methanogenesis and therefore generate comparable pH increases and bicarbonate shifts, likewise produced no precipitates despite incubation periods of 800 and 900 days respectively. The thermodynamic drive for HAp precipitation is therefore present in all biotic conditions, yet only *M.* AMB_03 overcomes the kinetic barrier to nucleation. This makes the *ppk1*-*ppx* genomic architecture, together with the co-occurring EPS/amyloid-associated genes unique to AMB_03 (Table 1; Section 4.5), the plausible genomic contributors to its biomineralization phenotype. With only three strains compared, the present dataset cannot fully separate the relative contributions of polyP cycling versus EPS-matrix composition; functional work (e.g., targeted gene disruption, heterologous expression, or additional *ppk1*-positive/negative strain pairs) would be needed to establish which, or both, is causal.

We therefore propose that elevated supersaturation driven by pH increase and EPS-mediated heterogeneous nucleation together overcome the kinetic barrier to HAp crystallisation. Blank solutions were also supersaturated with respect to calcite (SI = +0.3 to +0.6, Table S5), yet no carbonate phases were observed in blank experiments. This matches the expected pattern of kinetic inhibition of calcite nucleation by dissolved phosphate and Mg^2+^ present in the growth medium and even micromolar orthophosphate concentrations is enough to block active calcite growth sites by surface adsorption (Dove and Hochella, 1993; Lin and Singer, 2006). The diversity of carbonate morphologies in these low-phosphate experiments, ranging from sub-500 nm rose-textured spheres to pointed ovals and dumbbell-arranged calcite spikes (Fig. 4e-f), likely reflects the interplay between EPS organic matrix templates and varying local supersaturation states (Dupraz et al. 2009; Mercedes-Martín et al. 2021). The distinct morphologies suggest that EPS provides heterogeneous nucleation surfaces with differing surface chemistries at different locations, directing carbonate crystal growth along multiple trajectories (Kawaguchi et al. 2002). This biologically induced morphological diversity contrasts with blank experiments, where EDS confirmed that C- and O-bearing signals represent evaporitic medium residues lacking Ca (Table S1) rather than precipitated mineral phases, confirming that genuine carbonate nucleation requires biological surfaces (Braissant et al. 2003). The distinct Fe behaviour (Fig. 7) mirrors mackinawite dissolution in abiotic systems, where slow FeS dissolution releases Fe^2+^ into solution as the sulfide pool is consumed over time (Rickard, 2006), and with biological Fe uptake in inoculated conditions, where Fe is required for [NiFe]-hydrogenase and methyl-CoM reductase, essential enzymes in hydrogenotrophic methanogenesis. Combined with negative vivianite saturation indices across all conditions (SI = −0.8 to −3.5, Table S5), this confirms the spherical precipitates are Ca-P rather than Fe-P phases. The geochemical evidence thus constrains the observed precipitates to Ca-P phases; the question of what drives their formation in AMB_03 but not in the two comparator strains tested here points toward the genomic architecture of this strain, though confirming this more broadly would require testing additional taxa, including other *ppk1*-containing lineages.

### 4.3 Genomic features

All living cells require machinery for inorganic phosphate uptake and basic phosphate handling, and predictably all three strains share these conserved components (Table 1). Polyphosphate metabolism is however fundamentally different: the ability to synthesize, accumulate, and hydrolyze polyP represents a specialized and phylogenetically restricted capacity that creates intracellular phosphate reservoirs and generates localized phosphate fluxes beyond what standard cellular P-handling can produce (Kornberg, 1995; Rao et al. 2009). *M. oryzae* and MAB1 completely lack this capacity, no *ppk1*, no *ppk2*, no canonical *ppx*, while *M.* AMB_03 encodes a complete *ppk1*-*ppx* cycle in a conserved operon.

The *ppk1* catalyzes processive phosphate transfer from ATP to polyP and is the principal enzyme for intracellular polyP synthesis in prokaryotes (Ahn and Kornberg, 1990; Kornberg, 1995). ppk1 transcript levels directly regulate polyP accumulation in methanogenic archaea (Paula et al., 2019), consistent with an active polyP synthesis capacity in AMB_03. Neither *M. oryzae* nor MAB1 encodes *ppk1* or a canonical *ppx*; both possess *SurE*-type proteins annotated with EC 3.6.1.11 that share functional overlap with PPX but belong to a distinct protein family, and their exopolyphosphatase activity has not been experimentally confirmed. The complete *ppk1*-*ppx* cycle exclusively in AMB_03 is therefore consistent with active intracellular phosphate buffering and may provide the phosphate flux necessary to drive local supersaturation and extracellular Ca-P precipitation under energy-limited conditions.

An alternative interpretation is that the pH increase driven by hydrogenotrophic methanogenesis consuming dissolved CO_2_ is sufficient alone to elevate HAp supersaturation and drive precipitation, without invoking polyP cycling specifically. This cannot be excluded on thermodynamic grounds, since both inoculated and blank solutions reached HAp supersaturation (Table S5). However, three observations collectively argue against passive pH-driven precipitation as the sole mechanism. First, blank solutions were supersaturated (SI = +1.2 to +5.4) yet produced no Ca-P precipitates over the entire experiment, demonstrating that supersaturation is not sufficient for precipitation. Second, the temporal onset of precipitation, absent during active weekly feeding and developing progressively only under energy limitation, mirrors the expected dynamics of *ppx*-mediated polyP hydrolysis under starvation (Rao and Kornberg, 1996), not the pH trajectory, which would be expected to correlate with methanogenic activity rather than intensify as activity declines. Third, neither *M. oryzae* nor MAB1 produced precipitates under identical conditions including identical pH buffering capacity, ruling out a purely pH-mediated mechanism. These observations indicate polyP cycling as a necessary contributor, acting together with pH increase and EPS-mediated nucleation, and pH increase by itself does not account for the pattern. Beyond ruling out pH increase as the sole driver, multiple non-mutually exclusive mechanisms can explain how polyP cycling specifically promotes HAp precipitation in AMB_03. Under energy starvation, *ppx*-mediated degradation of accumulated polyP regenerates ATP and releases free orthophosphate (Rao and Kornberg, 1996) that diffuses to the extracellular space. Prolonged starvation may additionally trigger cell lysis, releasing intracellular polyP directly into the medium where dissolved Ca^2+^ and elevated pH create supersaturating conditions for HAp (Rao et al., 2009). EPS-bound polyP chains may further amplify local phosphate concentrations at the cell surface, as polyP has been shown to associate with extracellular matrices in phosphate-scavenging contexts (Rao et al., 2009). These mechanisms collectively predict precipitation delayed relative to growth onset and intensifying under energy limitation, precisely the pattern observed: no precipitates at week 6 during active feeding, developing progressively as feeding became infrequent, and most pronounced after 45-49 weeks of energy limitation. This temporal coupling distinguishes biologically induced mineralization from passive abiotic precipitation. The complete absence of precipitation in both *ppk1*-lacking comparators under identical conditions (section 2.1), combined with the temporal coupling to energy limitation, provides substantial evidence for a functional role of the polyphosphate synthesis machinery in this biomineralization phenotype. The coexistence of intracellular polyP cycling (Paula et al., 2019) and extracellular Ca-P precipitation suggests AMB_03 employs condition-dependent strategies across intracellular and extracellular compartments, linking mineral precipitation to active phosphorus metabolism rather than passive adsorption.

In the natural P-poor serpentinizing environment from where AMB_03 was isolated, this biomineralization capacity would be geochemically silent but could activate during episodic phosphate enrichment from hydrothermal fluid pulses or weathering inputs. Reduced phosphorus species such as phosphite may represent transient bioavailable P sources in such systems (Pasek et al., 2022), though much of the phosphorus inventory is ultimately predicted to be sequestered as Ca-P phases. The combined capacity for polyP metabolism and Ca-P biomineralization provides a plausible mechanism for local phosphate enrichment and stabilization: by coupling intracellular sequestration with extracellular authigenic precipitation, AMB_03 could transform a diffuse resource into a concentrated biological asset. The formation of labile or amorphous Ca-P phases at microbial surfaces may serve as a phosphate buffer against P-depleted bulk fluids, potentially mitigating the phosphate problem by maintaining high local P-density in the immediate vicinity of the biomass despite global geochemical scarcity.

This contrast in ecological context may help explain why polyP cycling is retained in AMB_03 but absent in its comparators, though not simply as a matter of phosphate-poor versus phosphate-rich habitats. *M. oryzae* was isolated from ricefield soil (Joulian et al., 2000), and *M. bourgensis* MAB1 has been recovered from biogas digesters (Manzoor et al. 2016); both settings are typically phosphate-replete and buffered by continuous agricultural or organic inputs, making phosphate supply comparatively stable rather than scarce. Serpentinizing systems such as Chimaera are, by contrast, chronically phosphate-poor at baseline but subject to episodic phosphate pulses from hydrothermal fluids or weathering (see above). Under a fluctuating regime of this kind, luxury polyP storage, accumulating phosphate when transiently available for use when it is not, would carry a selective advantage that a consistently phosphate-rich or consistently phosphate-poor environment would not provide. The absence of *ppk1-ppx* in both comparator strains is therefore consistent with polyP cycling capacity tracking exposure to fluctuating, rather than simply scarce, phosphate availability.

### 4.4 Phylogenetic distribution of polyphosphate metabolism genes across *Methanobacteriota*

A phylogenomic analysis of 320 representative *Methanobacteriota* genomes (Fig.8) places *M.* AMB_03 within the *Methanobacteriales* and reveals that the capacity for complete polyP cycling is far from universal in this phylum, establishing an evolutionary context for the strain-level differences observed here. As established in Section 3.6.3, complete polyP cycling capacity is uncommon within Methanobacteriota (∼25% of genomes; Table S6a), providing phylum-wide context for the strain-level contrast observed in AMB_03 versus its comparators. The predominance of *ppk1* over *ppk2* suggests the former is the primary polyphosphate synthesis pathway in this phylum. Intriguingly, 26% of *ppx*-containing genomes (28/107) lacked a corresponding synthesis gene, potentially indicating either incomplete pathway assembly or metabolic specialisation for polyphosphate scavenging. Organisms encoding *ppx* without a cognate synthesis gene may represent polyphosphate scavengers capable of utilising environmental or community-derived polyP as a phosphate source, analogous to extracellular polyP utilisation described in some bacteria (Rao et al., 2009). Alternatively, this pattern may reflect evolutionary gene loss of synthesis capacity while retaining degradation for phosphate mobilisation under starvation. Horizontal gene transfer of individual polyP pathway components, documented across prokaryotes (Kornberg, 1995), could also produce the mosaic distribution observed across *Methanobacteriota* genera. The finding that ∼25% of Methanobacteriota genomes possess complete polyP cycling capacity implies that this biomineralization potential is not restricted to serpentinizing environments but extends to the marine sedimentary methanic zone globally. Given the widespread distribution of Methanobacteriota in marine and freshwater sediments, polyP-capable methanogens may represent a previously unrecognised contributor to authigenic phosphate mineral formation below the SMT, complementing rather than replacing the established anaerobic methane oxidation pathway. Testing this at broader scale, beyond the GTDB species representatives used here is now feasible using large, quality-controlled archaeal genome collections such as ArchaeaHQ (Bespiatykh and Leao, 2026), whose ecologically-categorized metagenome-assembled genomes would allow polyP-cycling capacity to be mapped directly onto habitat. The 26% of *ppx*-containing genomes that lack a cognate synthesis gene are particularly intriguing in this context: if these organisms function as polyP scavengers, acquiring community-derived polyP as a phosphorus source rather than synthesizing it de novo, they could modulate the local phosphate inventory in sediments without being detectable by ppk1 presence alone. Calcium-phosphate biomineralization joins a small but growing set of biogeochemical processes whose capacity can be read directly from genomes through curated functional gene modules, as established for iron cycling (FeGenie; Garber et al., 2020) and hydrocarbon degradation (CANT-HYD; Khot et al., 2022). Our results suggest that for microbially mediated apatite formation the diagnostic unit is not a single gene but the co-occurrence of an ion-concentrating module (the *ppk1*–*ppx* cycle), consistent with the recognised role of microbial polyphosphate as a precursor of authigenic apatite in marine sediments (Diaz et al., 2008).

Consistent with the strain-level annotation (Section 3.6.2), AMB_03’s *ppk1* is immediately upstream of *ppx*/*GppA* (AMB03_00141), recapitulating the *ppk*–*ppx* gene arrangement common in prokaryotic polyphosphate metabolism. The co-localisation of these opposing enzymatic activities suggests that polyphosphate turnover is encoded in close genomic proximity, pointing to tight local regulation of the polyphosphate pool. This *ppk1*–*ppx* co-localisation is not unique to AMB_03: among the Methanobacteriota genomes encoding a complete pathway, the two genes are co-located on the same strand in AMB_03’s nearest relatives (Methanobacterium_D) and in cultured Methanosphaera references (*M*. *stadtmanae*, *M*. *massiliense)*, with *ppk1* amino-acid identity tracking phylogenetic distance (Table S7). Conservation of this arrangement across the clade indicates an ancestral *ppk1*–*ppx* pairing and supports interpreting the two adjacent *ppk1* ORFs in AMB_03 as a single gene split by an assembly artifact (Section 3.6.2) rather than a duplication. Matching the genomic analysis in Section 3.6.2, neither *M. oryzae* nor MAB1 harboured polyphosphate metabolism genes. Computational screening of the AMB_03 proteome identified candidate amyloid-forming proteins that could contribute to the physicochemical properties of the extracellular matrix.

### 4.5 Amyloid-like EPS components as candidate nucleation scaffolds

Beyond polyphosphate metabolism, the molecular basis of EPS-mediated nucleation in AMB_03 may involve an additional layer of structural biology. Recent studies have demonstrated that archaea produce functional amyloids contributing to extracellular structures. In methanogenic archaea, proteinaceous sheaths surrounding cells have been shown to exhibit amyloid-like properties, which explain their remarkable mechanical stability and resistance to degradation (Christensen et al., 2018; Dueholm et al., 2015). Archaea biofilms can also contain extracellular matrices composed of polysaccharides, extracellular DNA, and amyloid-like proteins (Chimileski et al., 2014), suggesting that amyloid assemblies may represent a conserved structural component of archaeal extracellular polymeric substances. More broadly, archaeal prion-like domains have been experimentally shown to form amyloid fibrils, supporting the use of computational approaches such as PLAAC to identify candidate amyloid-forming proteins in archaeal genomes (Zajkowski et al., 2021).

In *M.* AMB_03, PLAAC identified 12 candidates with prion-like domains (COREscore > 0), including proteins associated with cell-envelope function, metal binding, and polysaccharide synthesis, alongside seven hypothetical proteins (Table S8). These candidates are relevant because amyloid assemblies are increasingly recognised not only as structural scaffolds, but also as chemically active surfaces capable of binding metal ions and, in some cases, displaying enzyme-like catalytic activity (Esposito et al., 2026). Such properties could be important in the context of extracellular phosphate biomineralisation. Amyloid-like proteins within the EPS may concentrate Ca^2+^ and phosphate at the cell surface, provide ordered heterogeneous nucleation templates, and potentially influence phosphate availability through localised catalytic reactions. Among these candidates, the pseudomurein-binding repeat protein (M_AMB03_00342) is notable: pseudomurein is the structural cell wall polymer of *Methanobacteriales*, and cell-wall-anchored repeat-domain proteins are well positioned to form extracellular templates for mineral nucleation, presenting ordered surfaces at the precise location where HAp spheres are observed growing on and around cells (Fig. 4c-d). The glycosyltransferase family 39 protein (M_AMB03_02928) likely contributes to the EPS polysaccharide backbone visible in SEM (Fig. 3e-f, Fig. S4), providing the organic matrix scaffold within which HAp nucleation occurs. The cupredoxin domain protein (M_AMB03_00895), with its metal-coordinating capacity, represents a candidate for localised Ca²⁺ concentration at the cell-mineral interface, analogous to metal-binding protein assemblies in vertebrate enamel biomineralisation (Carneiro et al., 2016).

Analogous principles have been demonstrated in calcium-based biomineralization systems outside the archaeal domain. In studies of vertebrate enamel formation and biomimetic mineralization, protein assemblies with amyloid-like or highly ordered structures have been shown to bind calcium ions and direct the nucleation and growth of mineral phases (Carneiro et al., 2016; Castellanos et al., 2018). Although these systems are phylogenetically distant from archaea, they illustrate general physicochemical mechanisms by which ordered protein scaffolds can control mineral deposition.

Although direct evidence for amyloid formation or catalytic activity in AMB_03 is currently lacking, the co-occurrence of EPS production, extracellular Ca-P precipitation, and predicted prion-like proteins suggests a plausible mechanistic link between amyloid-like matrix components and the spatial control of mineralisation. Thus, amyloid-like proteins may contribute not only to extracellular aggregation and matrix stability, but also to the localisation and morphology of HAp precipitation.

### 4.6 Implications for phosphatization taphonomy

The methanogenic zone discussed here as a modern phosphate sink (Sections 1.3–1.4) lies below the sulfate-methane transition zone classically invoked in models of authigenic Ca-P formation and soft-tissue phosphatization. Microbially mediated apatite replacement of decaying soft tissue, best documented in Ediacaran-Cambrian phosphorite deposits such as the Doushantuo Formation (Xiao et al., 1998), has generally been attributed to sulfate-reducing bacteria and the decay of associated microbial mats, which locally raise phosphate and alkalinity at or within centimetres of the sediment-water interface (Wilby et al., 1996; Wilby and Briggs, 1997; Briggs, 2003). Our results raise the possibility that hydrogenotrophic methanogens with an active *ppk1-ppx* cycle represent an additional, previously unconsidered biological source of authigenic Ca-P relevant to phosphatization taphonomy, distinct in both setting and mechanism from the SMT-proximal, sulfate-reducer-driven pathway. We do not propose that methanogen-mediated precipitation explains phosphatization broadly, since classic settings for exceptional soft-tissue preservation are shallow marine and SMT-proximal rather than the deeper methanogenic zone occupied by AMB_03 and its relatives; rather, we suggest it broadens the set of microbial mechanisms capable of generating the localized Ca-P supersaturation that phosphatization requires, a possibility that could be tested where methanogenic zones intersect phosphatized fossil-bearing horizons in the rock record.

## 5. Conclusions

This study establishes three key findings: (1) among the strains tested, only *Methanobacterium* AMB_03 precipitates extracellular spheres of poorly crystalline, carbonate-substituted HAp, which to our knowledge is the first demonstration of HAp biomineralization linked to polyphosphate metabolism in a hydrogenotrophic methanogen; (2) the presence of a complete *ppk1*-*ppx* polyphosphate cycling system is strongly associated with this biomineralization capacity: a Raman band in the polyP window was detected in AMB_03 but not in *M. oryzae*, and both comparator strains (*M. oryzae* and *M. bourgensis* MAB1), which lack these genes, failed to mineralize despite comparable supersaturation, indicating that AMB_03, possibly through its EPS matrix, overcomes a kinetic barrier to nucleation; establishing causality will require functional validation beyond the correlative genomic comparison presented here; and (3) the capacity for complete polyP cycling is phylogenetically uncommon within *Methanobacteriota*, present in only ∼25% of 320 surveyed genomes, with AMB_03 additionally encoding candidate amyloid-like proteins that may contribute to EPS structure and heterogeneous nucleation. These results expand our understanding of microbial phosphorus cycling and suggest that polyphosphate-capable methanogens may create localized phosphate-rich niches in sedimentary environments, extending the known biological and geochemical drivers of authigenic Ca-P and Fe-P formation below the sulfate-methane transition (Sections 1.3–1.4) to include methanogen-derived polyP cycling as an additional, previously unrecognised pathway. The association of a complete *ppk1-ppx* polyphosphate cycle with biomineralization capacity further suggests a practical application: genomic screening for polyP synthesis capacity could identify methanogens with potential utility in phosphate recovery from engineered systems. Because sedimentary phosphorus burial is itself a primary control on ocean oxygenation over geological time (Section 1.2; Watson et al., 2017), a previously unrecognised biological contributor to that burial flux is relevant to how the marine phosphorus cycle, and its coupling to ocean redox history, is reconstructed. Given that the capacity for polyP cycling is scattered across several Methanobacteriota lineages (Section 4.4) rather than confined to serpentinizing settings, this contribution may recur wherever anoxic, methanogenic sediments have existed through Earth history, leaving a testable signature in phosphatized fossil-bearing horizons (Section 4.6).

## Acknowledgements

This study was supported by the Swedish research council (VR) project to A.N. (#2017-05018). T.Z. acknowledges support from National Science Centre, Poland (NCN), through OPUS 29 grant 2025/57/B/NZ8/03307. T.B. and O.T. thank the Olle Engkvist Stiftelse (project 235-0413) and the Wallenberg Initiative Materials Science for Sustainability (WISE), funded by the Knut and Alice Wallenberg Foundation, for their support.

## Data availability

The genome sequence of Methanobacterium AMB_03 is being deposited in NCBI GenBank; accession numbers will be provided in a revised version of this preprint. Raw gas chromatography, ICP-OES, Raman and XRD data, and the phylogenomic tree and gene-annotation files, are available from the corresponding author upon reasonable request.

## Author Contributions

A. Neubeck conceived and designed the study, performed laboratory work and analyses, wrote the manuscript, and acquired funding. This study builds in part on data originally generated during F.A. Robin-Champigneul’s MSc thesis project. This included laboratory work and analyses as well as writing. A. Singh performed most of the comparative genomic analysis. S. Sharma performed genomic and phylogenetic analyses and contributed with writing. O. Tkachenko performed mineralogical analyses, including XRD. J. Thyr contributed to spectroscopic analysis. S. Sidiropoulos contributed samples used in this study. A. Schnürer provided laboratory access and co-isolated *M. AMB_03*. T. Zajkowski performed the prediction of amyloid-forming (prion-like) proteins. T. Budnyak contributed to reviewing and editing the manuscript. All authors reviewed and approved the final manuscript.

## References

Ahn, K., Kornberg, A., 1990. Polyphosphate kinase from Escherichia coli. Purification and demonstration of a phosphoenzyme intermediate. J. Biol. Chem. 265, 11734–11739.

Araujo, J.C., Téran, F.C., Oliveira, R.A., Nour, E.A.A., Montenegro, M.A.P., Campos, J.R., Vazoller, R.F., 2003. Comparison of hexamethyldisilazane and critical point drying treatments for SEM analysis of anaerobic biofilms and granular sludge. J. Electron Microsc. (Tokyo) 52, 429–433. 10.1093/jmicro/52.4.429

Awonusi, A., Morris, M.D., Tecklenburg, M.M.J., 2007. Carbonate assignment and calibration in the Raman spectrum of apatite. Calcified Tissue International 81, 46–52.

Berner, R.A., 1990. Diagenesis of phosphorus in sediments from non-upwelling areas. Phosphate Depos. World Vol 3 Neogene Mod. Phosphorites 27–32.

Bespiatykh, D., Leão, P., 2026. ArchaeaHQ: A Curated Reference Database of Archaeal Genomes. bioRxiv. 10.64898/2026.06.02.729493

Braissant, O., Cailleau, G., Dupraz, C., Verrecchia, E.P., 2003. Bacterially Induced Mineralization of Calcium Carbonate in Terrestrial Environments: The Role of Exopolysaccharides and Amino Acids. Journal of Sedimentary Research, 73(3), 485–490. 10.1306/111302730485

Bray, J.T., Bricker, O.P., Troup, B.N., 1973. Phosphate in Interstitial Waters of Anoxic Sediments: Oxidation Effects during Sampling Procedure. Science 180, 1362–1364. 10.1126/science.180.4093.1362

Briggs, D.E.G., 2003. The role of decay and mineralization in the preservation of soft-bodied fossils. Annual Review of Earth and Planetary Sciences 31, 275–301.

Burns, S.J., 1997. Early diagenesis in Amazon fan sediments. Early Diagenesis Amaz. Fan Sediments 155, 497–504.

Capella-Gutiérrez, S., Silla-Martínez, J.M., Gabaldón, T., 2009. trimAl: a tool for automated alignment trimming in large-scale phylogenetic analyses. Bioinformatics 25, 1972–1973. 10.1093/bioinformatics/btp348

Cantalapiedra, C.P., Hernández-Plaza, A., Letunic, I., Bork, P., Huerta-Cepas, J., 2021. eggNOG-mapper v2: functional annotation, orthology assignments, and domain prediction at the metagenomic scale. Mol. Biol. Evol. 38, 5825–5829. 10.1093/molbev/msab293

Carneiro, K.M.M., Zhai, H., Zhu, L., Horst, J.A., Sitlin, M., Nguyen, M., Wagner, M., Simpliciano, C., Milder, M., Chen, C.-L., Ashby, P., Bonde, J., Li, W., Habelitz, S., 2016. Amyloid-like ribbons of amelogenins in enamel mineralization. Sci. Rep. 6, 23105. 10.1038/srep23105

Castellanos, M., Torres-Pardo, A., Rodríguez-Pérez, R., Gasset, M., 2018. Amyloid Assembly Endows Gad m 1 with Biomineralization Properties. Biomolecules 8, 13. 10.3390/biom8010013

Chimileski, S., Franklin, M.J., Papke, R.T., 2014. Biofilms formed by the archaeon Haloferax volcanii exhibit cellular differentiation and social motility, and facilitate horizontal gene transfer. BMC Biol. 12, 65. 10.1186/s12915-014-0065-5

Christensen, L.F.B., Hansen, L.M., Finster, K., Christiansen, G., Nielsen, P.H., Otzen, D.E., Dueholm, M.S., 2018. The Sheaths of Methanospirillum Are Made of a New Type of Amyloid Protein. Front. Microbiol. 9, 2729. 10.3389/fmicb.2018.02729

Czamara, K. et al. (2015). Raman spectroscopy of lipids: a review. Journal of Raman Spectroscopy, 46(1), 4–20. doi:10.1002/jrs.4607

De Gelder, J., De Gussem, K., Vandenabeele, P., Moens, L., 2007. Reference database of Raman spectra of biological molecules. J. Raman Spectrosc. 38, 1133–1147. 10.1002/jrs.1734

Delaney, M.L., 1998. Phosphorus accumulation in marine sediments and the oceanic phosphorus cycle. Glob. Biogeochem. Cycles 12, 563–572. 10.1029/98GB02263

Diaz, J., Ingall, E., Benitez-Nelson, C., Paterson, D., de Jonge, M.D., McNulty, I., Brandes, J.A., 2008. Marine Polyphosphate: A Key Player in Geologic Phosphorus Sequestration. Science 320, 652–655. 10.1126/science.1151751

Dove, P.M., Hochella, M.F., 1993. Calcite precipitation mechanisms and inhibition by orthophosphate: In situ observations by Scanning Force Microscopy. Geochim. Cosmochim. Acta 57, 705–714. 10.1016/0016-7037(93)90381-6

Dueholm, M.S., Larsen, P., Finster, K., Stenvang, M.R., Christiansen, G., Vad, B.S., Bøggild, A., Otzen, D.E., Nielsen, P.H., 2015. The Tubular Sheaths Encasing Methanosaeta Thermophila Filaments Are Functional Amyloids. J. Biol. Chem. 290, 20590–20600. 10.1074/jbc.M115.654780

Dupraz, C., Reid, R., Braissant, O., Decho, A., Norman, R., Visscher, P., 2009. Processes of carbonate precipitation in modern microbial mats. Earth-Sci. Rev. 96, 141–162. 10.1016/j.earscirev.2008.10.005

Eddy, S.R., 2011. Accelerated Profile HMM Searches. PLoS Comput. Biol. 7, e1002195. 10.1371/journal.pcbi.1002195

Edgar, R.C., 2004. MUSCLE: a multiple sequence alignment method with reduced time and space complexity. BMC Bioinformatics 5, 113. 10.1186/1471-2105-5-113

Egger, M., Jilbert, T., Behrends, T., Rivard, C., Slomp, C.P., 2015. Vivianite is a major sink for phosphorus in methanogenic coastal surface sediments. Geochim. Cosmochim. Acta 169, 217–235. 10.1016/j.gca.2015.09.012

Egger, M., Kraal, P., Jilbert, T., Sulu-Gambari, F., Sapart, C.J., Röckmann, T., Slomp, C.P., 2016. Anaerobic oxidation of methane alters sediment records of sulfur, iron and phosphorus in the Black Sea. Biogeosciences 13, 5333–5355. 10.5194/bg-13-5333-2016

Ensikat, H.-J., Geisler, T., Weigend, M., 2016. A first report of hydroxylated apatite as structural biomineral in Loasaceae – plants’ teeth against herbivores. Sci. Rep. 6, 26073. 10.1038/srep26073

Esposito, A., Leone, L., Nastri, F., Lombardi, A., 2026. Catalytic Amyloids: Turning Fibrils Into Biocatalysts. Chem. Eur. J. 32, e202503095. 10.1002/chem.202503095

Fagel, N., Alleman, L.Y., Granina, L., Hatert, F., Thamo-Bozso, E., Cloots, R., André, L., 2005. Vivianite formation and distribution in Lake Baikal sediments. Glob. Planet. Change, Progress towards reconstructing past climate in Central Eurasia, with special emphasis on Lake Baikal 46, 315–336. 10.1016/j.gloplacha.2004.09.022

Filippelli, G.M., 1997. Controls on phosphorus concentration and accumulation in oceanic sediments. Mar. Geol, Phosphorus Cycling and Sedimentation in Marine and Freshwater Systems 139, 231–240. 10.1016/S0025-3227(96)00113-2

Garber, A.I., Nealson, K.H., Okamoto, A., McAllister, S.M., Chan, C.S., Barco, R.A., Merino, N., 2020. FeGenie: A Comprehensive Tool for the Identification of Iron Genes and Iron Gene Neighborhoods in Genome and Metagenome Assemblies. Front. Microbiol. 11, 37. 10.3389/fmicb.2020.00037

Gemeri, D., Živković, A., Lukačević, I., Bahmann, H., King, H.E., 2023. Density functional theory demonstrates orientation effects in the Raman spectra of hydroxy-and carbonated apatite. J. Raman Spectrosc. 54, 159–170. 10.1002/jrs.6465

Graf, D.L., 1961. Crystallographic tables for the rhombohedral carbonates. Am. Mineral. 46, 1283–1316.

Gu, W., Müller, A.L., Deutzmann, J.S., Williamson, J.R., Spormann, A.M., 2022. Growth rate-dependent coordination of catabolism and anabolism in the archaeon Methanococcus maripaludis under phosphate limitation. ISME J. 16, 2313–2319. 10.1038/s41396-022-01278-9

Hendra, P.J., Maddams, W.F., Royaud, I.A.M., Willis, H.A. & Zichy, V. (1990). Spectrochimica Acta Part A, 46, 747–756.

Hsu, T.-W., Jiang, W.-T., Wang, Y., 2014. Authigenesis of vivianite as influenced by methane-induced sulfidization in cold-seep sediments off southwestern Taiwan. J. Asian Earth Sci. 89, 88–97. 10.1016/j.jseaes.2014.03.027

Huerta-Cepas, J., Serra, F., Bork, P., 2016. ETE 3: Reconstruction, Analysis, and Visualization of Phylogenomic Data. Mol. Biol. Evol. 33, 1635–1638. 10.1093/molbev/msw046

Huerta-Cepas, J., Szklarczyk, D., Heller, D., Hernández-Plaza, A., Forslund, S.K., Cook, H., Mende, D.R., Letunic, I., Rattei, T., Jensen, L.J., von Mering, C., Bork, P., 2019. eggNOG 5.0: a hierarchical, functionally and phylogenetically annotated orthology resource based on 5090 organisms and 2502 viruses. Nucleic Acids Res. 47, D309–D314. 10.1093/nar/gky1085

Hyatt, D., Chen, G.-L., LoCascio, P.F., Land, M.L., Larimer, F.W., Hauser, L.J., 2010. Prodigal: prokaryotic gene recognition and translation initiation site identification. BMC Bioinformatics 11, 119. 10.1186/1471-2105-11-119

Inskeep, W., Silvertooth, J., 1988. Kinetjcs of hydroxyapatite precipitation at pH 7.4 to 8.4. Geochim. Cosmochim. Acta 52, 1883–1893.

Joulian, C., Patel, B.K.C., Ollivier, B., Garcia, J.-L., Roger, P.A., 2000. *Methanobacterium oryzae* sp. nov., a novel methanogenic rod isolated from a Philippines ricefield. International Journal of Systematic and Evolutionary Microbiology 50, 525–528.

Kawaguchi, T., Decho, A.W., 2002. A laboratory investigation of cyanobacterial extracellular polymeric secretions (EPS) in influencing CaCO₃ polymorphism. Journal of Crystal Growth, 240(1–2), 230–235.

Kay, M.I., Young, R.A., Posner, A.S., 1964. Crystal structure of hydroxyapatite. Nature 204, 1050–1052. 10.1038/2041050a0

Khot, V., Zorz, J., Gittins, D.A., Chakraborty, A., Bell, E., Bautista, M.A., Paquette, A.J., Hawley, A.K., Novotnik, B., Hubert, C.R.J., 2022. CANT-HYD: A Curated Database of Phylogeny-Derived Hidden Markov Models for Annotation of Marker Genes Involved in Hydrocarbon Degradation. Front. Microbiol. 12, 764058. 10.3389/fmicb.2021.764058

Kidder, D.L., Worsley, T.R., 2010. Phanerozoic Large Igneous Provinces (LIPs), HEATT (Haline Euxinic Acidic Thermal Transgression) episodes, and mass extinctions. Palaeogeogr. Palaeoclimatol. Palaeoecol. 295, 162–191. 10.1016/j.palaeo.2010.05.036

Kornberg, A., 1995. Inorganic polyphosphate: toward making a forgotten polymer unforgettable. J. Bacteriol. 177, 491–496. 10.1128/jb.177.3.491-496.1995

Lancaster, A.K., Nutter-Upham, A., Lindquist, S., King, O.D., 2014. PLAAC: a web and command-line application to identify proteins with prion-like amino acid composition. Bioinformatics 30, 2501–2502. 10.1093/bioinformatics/btu310

Larralde, M., 2022. Pyrodigal: Python bindings and interface to Prodigal, an efficient method for gene prediction in prokaryotes. J. Open Source Softw. 7, 4296. 10.21105/joss.04296

Lee, M.D., 2019. GToTree: a user-friendly workflow for phylogenomics. Bioinformatics 35, 4162–4164. 10.1093/bioinformatics/btz188

Letunic, I., Bork, P., 2007. Interactive Tree Of Life (iTOL): an online tool for phylogenetic tree display and annotation. Bioinformatics 23, 127–128. 10.1093/bioinformatics/btl529

Li, W., Joshi, S.R., Hou, G., Burdige, D.J., Sparks, D.L., Jaisi, D.P., 2015. Characterizing Phosphorus Speciation of Chesapeake Bay Sediments Using Chemical Extraction, 31P NMR, and X-ray Absorption Fine Structure Spectroscopy. Environ. Sci. Technol. 49, 203–211. 10.1021/es504648d

Lin, Y.-P., Singer, P.C., 2006. Inhibition of calcite precipitation by orthophosphate: Speciation and thermodynamic considerations. Geochim. Cosmochim. Acta 70, 2530–2539. 10.1016/j.gca.2006.03.002

Majed, N., Matthäus, C., Diem, M., Gu, A.Z., 2009. Evaluation of Intracellular Polyphosphate Dynamics in Enhanced Biological Phosphorus Removal Process using Raman Microscopy. Environ. Sci. Technol. 43(14), 5436–5442. 10.1021/es900251n

Manzoor, S., Schnürer, A., Bongcam-Rudloff, E., Müller, B., 2016. Complete genome sequence of *Methanoculleus bourgensis* strain MAB1, the syntrophic partner of mesophilic acetate-oxidising bacteria (SAOB). Standards in Genomic Sciences 11, 80. 10.1186/s40793-016-0199-x

Mercedes-Martín, R., Rogerson, M., Prior, T.J., Brasier, A.T., Reijmer, J.J.G., Billing, I., Matthews, A., Love, T., Lepley, S., Pedley, M. (2021). Towards a morphology diagram for terrestrial carbonates: evaluating the impact of carbonate supersaturation and alginic acid in calcite precipitate morphology. Geochimica et Cosmochimica Acta, 306, 340–361. 10.1016/j.gca.2021.04.010

Metz R, Kumar N, Schenkeveld WDC, Kraemer SM. Rates and Mechanism of Vivianite Dissolution under Anoxic Conditions. Environ Sci Technol. 2023 Nov 14;57(45):17266–17277. doi: 10.1021/acs.est.3c04474. Epub 2023 Nov 4. PMID: 37924285; PMCID: PMC10653223.

Miller, J.V. & Bartick, E.G. (2001). Forensic analysis of single fibers by Raman spectroscopy. Applied Spectroscopy, 55(12), 1729–1732.

Mänd, K., Kirsimäe, K., Lepland, A., Crosby, C.H., Bailey, J.V., Konhauser, K.O., Wirth, R., Schreiber, A., Lumiste, K., 2018. Authigenesis of biomorphic apatite particles from Benguela upwelling zone sediments off Namibia: The role of organic matter in sedimentary apatite nucleation and growth. Geobiology 16, 640–658. 10.1111/gbi.12309

März, C., Hoffmann, J., Bleil, U., de Lange, G.J., Kasten, S., 2008. Diagenetic changes of magnetic and geochemical signals by anaerobic methane oxidation in sediments of the Zambezi deep-sea fan (SW Indian Ocean). Mar. Geol. 255, 118–130. 10.1016/j.margeo.2008.05.013

März, C., Riedinger, N., Sena, C., Kasten, S., 2018. Phosphorus dynamics around the sulphate-methane transition in continental margin sediments: Authigenic apatite and Fe(II) phosphates. Mar. Geol. 404, 84–96. 10.1016/j.margeo.2018.07.010

Medvedeff, C.A., Inglett, K.S., Inglett, P.W., 2014. Evaluation of direct and indirect phosphorus limitation of methanogenic pathways in a calcareous subtropical wetland soil. Soil Biol. Biochem. 69, 343–345. 10.1016/j.soilbio.2013.11.018

Neubeck, A., Callac, N., Isaksson, S., Schnürer, A., 2025. Growth of hydrogenotrophic methanogen Methanoculleus bourgensis MAB1 in the presence of dunite. Anaerobe 92, 102945. 10.1016/j.anaerobe.2025.102945

Neubeck, A., Sun, L., Müller, B., Ivarsson, M., Hosgörmez, H., Özcan, D., Broman, C., Schnürer, A., 2017. Microbial Community Structure in a Serpentine-Hosted Abiotic Gas Seepage at the Chimaera Ophiolite, Turkey. Appl. Environ. Microbiol. 83, e03430–16. 10.1128/AEM.03430-16

Oxford Nanopore Technologies. Medaka: sequence correction tool. https://github.com/nanoporetech/medaka (accessed 2026).

Parkhurst, D.L., Appelo, C. a. J., 2013. Description of input and examples for PHREEQC version 3: A computer program for speciation, batch-reaction, one-dimensional transport, and inverse geochemical calculations (No. 6-A43), Techniques and Methods. U.S. Geological Survey. 10.3133/tm6A43

Pasek, M.A., Omran, A., Feng, T., Gull, M., Lang, C., Abbatiello, J., Garong, L., Johnston, R., Ryan, J., Abbott-Lyon, H., 2022. Serpentinization as a route to liberating phosphorus on habitable worlds. Geochim. Cosmochim. Acta 336, 332–340. 10.1016/j.gca.2022.09.027

Passey, S., Pellegrin, S., Mellor, H., 2007. Scanning electron microscopy of cell surface morphology. Curr. Protoc. Cell Biol. Chapter 4, Unit4.17. 10.1002/0471143030.cb0417s37

Paula, F.S., Chin, J.P., Schnürer, A., Müller, B., Manesiotis, P., Waters, N., Macintosh, K.A., Quinn, J.P., Connolly, J., Abram, F., McGrath, J.W., O’Flaherty, V., 2019. The potential for polyphosphate metabolism in Archaea and anaerobic polyphosphate formation in Methanosarcina mazei. Sci. Rep. 9, 17101. 10.1038/s41598-019-53168-4

Pezzotti, G. (2021). Raman spectroscopy in cell biology and microbiology. Journal of Raman Spectroscopy, 52(12), 2348–2443. doi:10.1002/jrs.6204

Price, M.N., Dehal, P.S., Arkin, A.P., 2010. FastTree 2--approximately maximum-likelihood trees for large alignments. PloS One 5, e9490. 10.1371/journal.pone.0009490

Racki, L.R., Freddolino, L., 2025. Polyphosphate: The “Dark Matter” of Bacterial Chromatin Structure. Mol. Microbiol. 123, 279–293. 10.1111/mmi.15350

Rao, N.N., Gómez-García, M.R., Kornberg, A., 2009. Inorganic polyphosphate: essential for growth and survival. Annu. Rev. Biochem. 78, 605–647. 10.1146/annurev.biochem.77.083007.093039

Rao, N.N., Kornberg, A., 1996. Inorganic polyphosphate supports resistance and survival of stationary-phase Escherichia coli. J. Bacteriol. 178, 1394–1400. 10.1128/jb.178.5.1394-1400.1996

Rickard, D., 2006. The solubility of FeS. Geochim. Cosmochim. Acta 70, 5779–5789. 10.1016/j.gca.2006.02.029

Rudnick, H., Hendrich, S., Pilatus, U., Blotevogel, K.-H., 1990. Phosphate accumulation and the occurrence of polyphosphates and cyclic 2,3-diphosphoglycerate in Methanosarcina frisia. Arch. Microbiol. 154, 584–588. 10.1007/BF00248840

Scherer, P., Lippert, H., Wolff, G., 1983. Composition of the major elements and trace elements of 10 methanogenic bacteria determined by inductively coupled plasma emission spectrometry. Biol. Trace Elem. Res. 5, 149–163. 10.1007/BF02916619

Schwengers, O., Jelonek, L., Dieckmann, M.A., Beyvers, S., Blom, J., Goesmann, A., 2021. Bakta: rapid and standardized annotation of bacterial genomes via alignment-free sequence identification. Microb. Genom. 7, 000685. 10.1099/mgen.0.000685 eggNOG-mapper

Seemann, T., 2014. Prokka: rapid prokaryotic genome annotation. Bioinformatics 30, 2068–2069. 10.1093/bioinformatics/btu153

Serrano, P., Hermelink, A., Lasch, P., de Vera, J.-P., König, N., Burckhardt, O., Wagner, D., 2015. Confocal Raman microspectroscopy reveals a convergence of the chemical composition in methanogenic archaea from a Siberian permafrost-affected soil. FEMS Microbiol. Ecol. 91(12), fiv126. 10.1093/femsec/fiv126

Slomp, C.P., Mort, H.P., Jilbert, T., Reed, D.C., Gustafsson, B.G., Wolthers, M., 2013. Coupled Dynamics of Iron and Phosphorus in Sediments of an Oligotrophic Coastal Basin and the Impact of Anaerobic Oxidation of Methane. PLOS ONE 8, e62386. 10.1371/journal.pone.0062386

Toso, D.B., Henstra, A.M., Gunsalus, R.P., Zhou, Z.H., 2011. Structural, mass and elemental analyses of storage granules in methanogenic archaeal cells. Environ. Microbiol. 13, 2587–2599. 10.1111/j.1462-2920.2011.02531.x

Vaser, R., Sović, I., Nagarajan, N., Šikić, M., 2017. Fast and accurate de novo genome assembly from long uncorrected reads. Genome Res. 27, 737–746. 10.1101/gr.214270.116

Watson, A.J., Lenton, T.M., Mills, B.J.W., 2017. Ocean deoxygenation, the global phosphorus cycle and the possibility of human-caused large-scale ocean anoxia. Philos. Trans. R. Soc. Math. Phys. Eng. Sci. 375, 20160318. 10.1098/rsta.2016.0318

Weiner, S., Addadi, L., 1997. Design strategies in mineralized biological materials. J. Mater. Chem. 7, 689–702. 10.1039/a604512j

Westerholm, M., Roos, S., Schnürer, A., 2010. Syntrophaceticus schinkii gen. nov., sp. nov., an anaerobic, syntrophic acetate-oxidizing bacterium isolated from a mesophilic anaerobic filter. FEMS Microbiology Letters 309, 100–104.10.1128/AEM.01637-12

Westerholm, M., Levén, L., Schnürer, A., 2012. Bioaugmentation of Syntrophic Acetate-Oxidizing Culture in Biogas Reactors Exposed to Increasing Levels of Ammonia. Appl. Environ. Microbiol. 78, 7619. 10.1128/AEM.01637-12

Wilby, P.R., Briggs, D.E.G., Bernier, P., Gaillard, C., 1996. Role of microbial mats in the fossilization of soft tissues. Geology 24, 787–790.

Wilby, P.R., Briggs, D.E.G., 1997. Taxonomic trends in the resolution of detail preserved in fossil phosphatized soft tissues. Geobios 30, 493–502.

Xiao, S., Zhang, Y., Knoll, A.H., 1998. Three-dimensional preservation of algae and animal embryos in a Neoproterozoic phosphorite. Nature 391, 553–558.

Zajkowski, T., Lee, M.D., Mondal, S.S., Carbajal, A., Dec, R., Brennock, P.D., Piast, R.W., Snyder, J.E., Bense, N.B., Dzwolak, W., Jarosz, D.F., Rothschild, L.J., 2021. The Hunt for Ancient Prions: Archaeal Prion-Like Domains Form Amyloid-Based Epigenetic Elements. Mol. Biol. Evol. 38, 2088–2103. 10.1093/molbev/msab010

Zajkowski, T., Lee, M.D., Sharma, S., Vallota-Eastman, A., Kuska, M., Malczewska, M., Rothschild, L.J., 2023. Conserved functions of prion candidates suggest a primeval role of protein self-templating. Proteins 91, 1298–1315. 10.1002/prot.26558

Zehnder, A.J.B., Huser, B.A., Brock, T.D., Wuhrmann, K., 1980. Characterization of an acetate-decarboxylating, non-hydrogen-oxidizing methane bacterium. Archives of Microbiology 124, 1–11. 10.1007/BF00407022

Zhuang, L., Tang, Z., Yu, Z., Li, J., Tang, J., 2019. Methanogenic Activity and Microbial Community Structure in Response to Different Mineralization Pathways of Ferrihydrite in Paddy Soil. Front. Earth Sci. 7. 10.3389/feart.2019.00325

