## Supplementary Material for "Polyphosphate metabolism genes are associated with hydroxyapatite biomineralization capacity in a novel hydrogenotrophic methanogen *Methanobacterium* AMB_03"

***Table S1.** Setup of first batch experiments.*

First batch

| *Flask no.* | *BM (mL)* | *C1 (mL)* | *C2 (mL)* | *[P] (mM)* | *Inoculation* |
|---|---|---|---|---|---|
| *1-4* | 18 | 1 | 1 | 0 | 1mL *M. AMB_03* |
| *5-8* | 18 | 1 | 1 | 0 | None |
| *9-12* | 18 | 1 | 1 | 0.1 | 1mL *M. AMB_03* |
| *13-16* | 18 | 1 | 1 | 0.1 | None |
| *17-20* | 18 | 1 | 1 | 0.5 | 1mL *M. AMB_03* |
| *21-24* | 18 | 1 | 1 | 0.5 | None |
| *25-28* | 18 | 1 | 1 | 1.0 | 1mL *M. AMB_03* |
| *29-32* | 18 | 1 | 1 | 1.0 | None |
| *33-36* | 18 | 1 | 1 | 3.0 | 1mL *M. AMB_03* |
| *37-40* | 18 | 1 | 1 | 3.0 | None |
| *41-44* | 18 | 1 | 1 | 5.8 (≈6) | 1mL *M. AMB_03* |
| *45-48* | ≤18* | 1 | 1 | 5.8 (≈6) | None |

* flask 48 has slightly less than 18 mL BM, exact amount uncertain.

Second batch

| Flask no. | BM (mL) | C1 (mL) | C2 (mL) | [P] (mM) | Inoculation |
|---|---|---|---|---|---|
| *49, 50* | 18 | 1 | 1 | 0 | 1mL *M. AMB_03* |
| *51, 52* | 18 | 1 | 1 | 0.1 | 1mL *M. AMB_03* |
| *53, 54* | 18 | 1 | 1 | 0.5 | 1mL *M. AMB_03* |
| *55, 56* | 18 | 1 | 1 | 1.0 | 1mL *M. AMB_03* |
| *57, 58* | 18 | 1 | 1 | 3.0 | 1mL *M. AMB_03* |
| *59, 60* | 18 | 1 | 1 | 5.8 (≈6) | 1mL *M. AMB_03* |

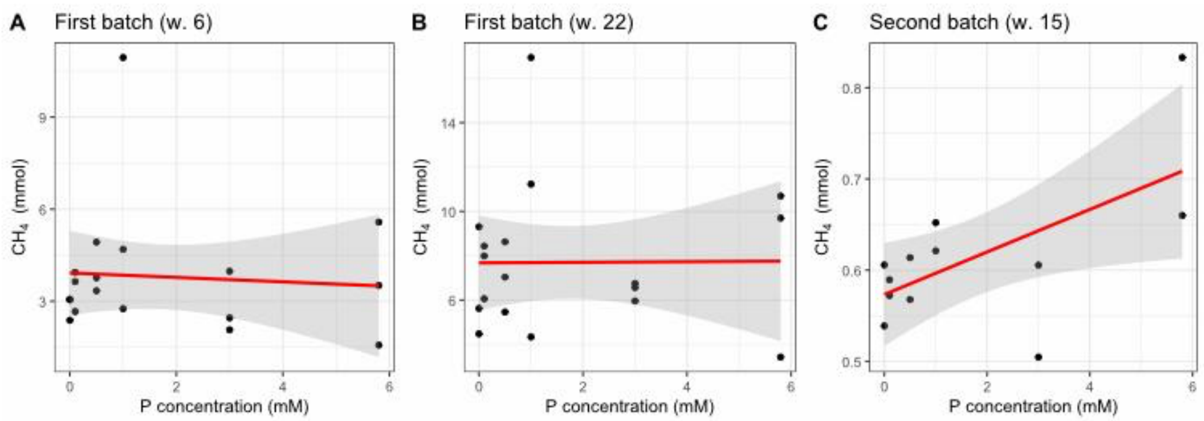


***Fig.S1.*** *Pearson correlation between cumulative $CH_4$ production and phosphate concentration across treatment levels. (A) batch 1 at week 6. (B) Batch 1 at week 22.(C) Batch 2 at week 15 under energy-limited conditions (single feeding). Points represent individual replicate bottles (n=18, Batch 1; n=12, Batch 2). Pearson correlation coefficient (r) and p-values are given in panel headers.*

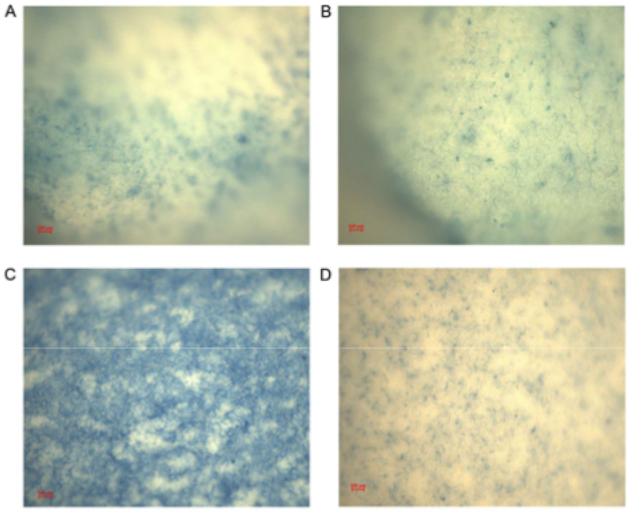


*Figure S2. Alcian Blue-stained filters observed under optical microscopy: inoculated (A) P = 0, (B) P = 3, (C) P =6, and blank (D) P = 6. Scale bars = 100 µm.*

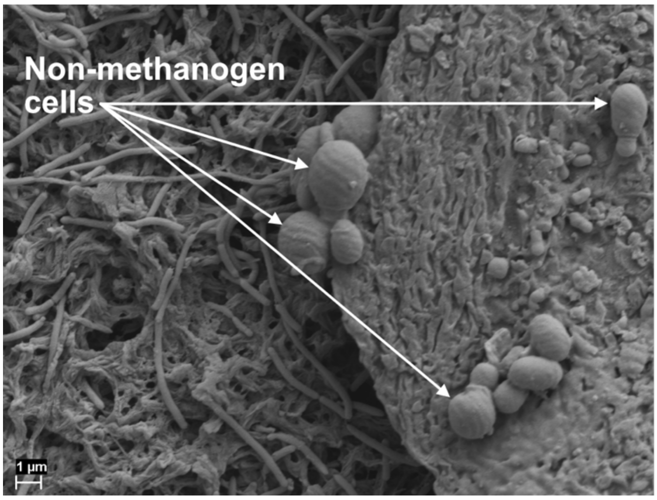


*Figure S3. SEM image with white arrows showing contaminant cells.*

***Table S2.*** EDS analyses of the HAp experiments.

| Experiment | Measurement | C (wt%) | O (wt%) | P (wt%) | Ca (wt%) | S (wt%) | Zn (wt%) | Fe (wt%) | Ca/P (atomic ratio) |
|---|---|---|---|---|---|---|---|---|---|
| Blank P0 | BLANK-P0-1 | 13.32 | 11.02 | 9.37 | — | 16.09 | 50.2 | — | — |
| Blank P0 | BLANK-P0-2 | 17.33 | 10.46 | — | — | 15.7 | 28.98 | — | — |
| Blank P0 | BLANK-P0-4 | 32.56 | 13.73 | 5.87 | — | 11.36 | 36.47 | — | — |
| Blank P3 | BLANK-P3 | 35.98 | 5.64 | — | — | 15.19 | 43.2 | — | — |
| P0 | P0-1-a (carbonate) | 18.11 | 47.65 | — | 28.2 | — | — | — | — |
| P0 | P0-1-b (cells) | 40.51 | 32.52 | — | 6.23 | — | — | 0.52 | — |

| | | | | | | | | | |
|---|---|---|---|---|---|---|---|---|---|
| P0 | P0-1-c (cells/matrix) | 28.8 | 33.79 | — | 5.77 | — | — | — | — |
| P0 | P0-3-a (cells/matrix) | 22.2 | 31.72 | 1.75 | 10.96 | — | — | — | 4.84 |
| P0 | P0-3-b (matrix) | 20.47 | 33.72 | — | 11.57 | — | — | — | — |
| P0.1 | P01-2 (cells/matrix) | 37.63 | 27.77 | 1.83 | 7.73 | — | — | 0.77 | 3.27 |
| P0.1 | P01-2-b (cells) | 37.93 | 28.15 | 2.01 | 8.05 | — | — | — | 3.10 |
| P0.1 | P01-2-c (cells/matrix) | 33.19 | 34.05 | — | 9.44 | — | — | — | — |
| P0.5 | P05-1 (cells) | 75.47 | 13.02 | 5.09 | — | — | — | — | — |
| P0.5 | P05-3-a (EPS/organic) | 80.51 | 17.92 | — | 0.63 | — | — | — | — |
| P0.5 | P05-5 (organic) | 77.67 | 15.55 | — | — | — | — | — | — |
| P1 | P1-2-a (cells/organic) | 86.24 | 12.37 | — | — | — | — | — | — |
| P1 | P1-2-b (sputter coat) | 74.19 | 7.03 | — | — | — | — | — | — |
| P1 | P1-3-a (organic) | 69.2 | 26.5 | — | — | — | — | — | — |
| P3 | P3-3 (cells) | 66.51 | 13.08 | 3.21 | — | — | — | — | — |
| P3 | P3bis-1 (organic) | 85.86 | 7.24 | — | — | — | — | — | — |
| P3 | P3bis-2 (cells) | 53.17 | 23.58 | — | — | — | — | — | — |
| P3 | P3bis-3 (HAp sphere) | 39.27 | 23.49 | 12.02 | 13.46 | — | — | — | 0.87 |
| P6 | P6-1 (HAp sphere) | 23.92 | 29.65 | 16.08 | 21.36 | — | — | — | 1.03 |
| P6 | P6-2 (HAp sphere) | 30.06 | 25.66 | 13.84 | 19.53 | — | — | — | 1.09 |
| P6 (Sotiris) | SOTIRIS-2 (cells) | 49.33 | 17.9 | 7.71 | 6.22 | — | — | 1.53 | 0.62 |
| P6 (Sotiris) | SOTIRIS-3 (HAp sphere) | 12.55 | 35.89 | 18.09 | 29.53 | — | — | — | 1.26 |
| P6 (Sotiris) | SOTIRIS-4 (cells/organic) | 62.78 | 19.34 | — | 1.56 | — | — | — | — |
| P6 (Sotiris) | SOTIRIS-8 (HAp+cells) | 28.17 | 33.09 | 12.02 | 11.62 | — | — | — | 0.75 |
| Field sediment (Chimaera) | Vitt-1 (carbonate) | 13.73 | 45.62 | — | 35.61 | — | — | — | — |
| Field sediment (Chimaera) | Vitt-1-2 (cells/matrix) | 40.74 | 32.65 | — | 7.62 | — | — | — | — |
| Field sediment (Chimaera) | Vitt-1-3 (matrix) | 35.15 | 39.26 | — | 5.82 | — | — | — | — |
| Field sediment (Chimaera) | Vitt-2-1 (serpentinite) | 9.64 | 32.65 | — | 17.2 | — | — | — | — |

| | | | | | | | | | |
|---|---|---|---|---|---|---|---|---|---|
| Field sediment (Chimaera) | Vitt-2-2 (serpentinite) | 11.7 | 34.47 | — | 15.49 | — | — | — | — |
| Field sediment (Chimaera) | Vitt-3 (carbonate) | 6.38 | 38.43 | — | 48.58 | — | — | — | — |

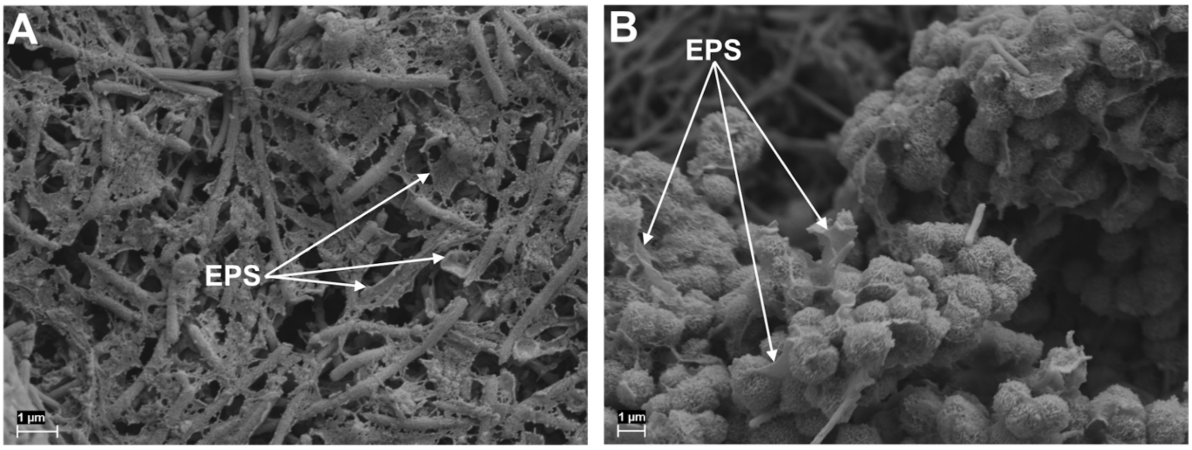


***Fig.S4.*** *SEM images with white arrows pointing towards EPS.*

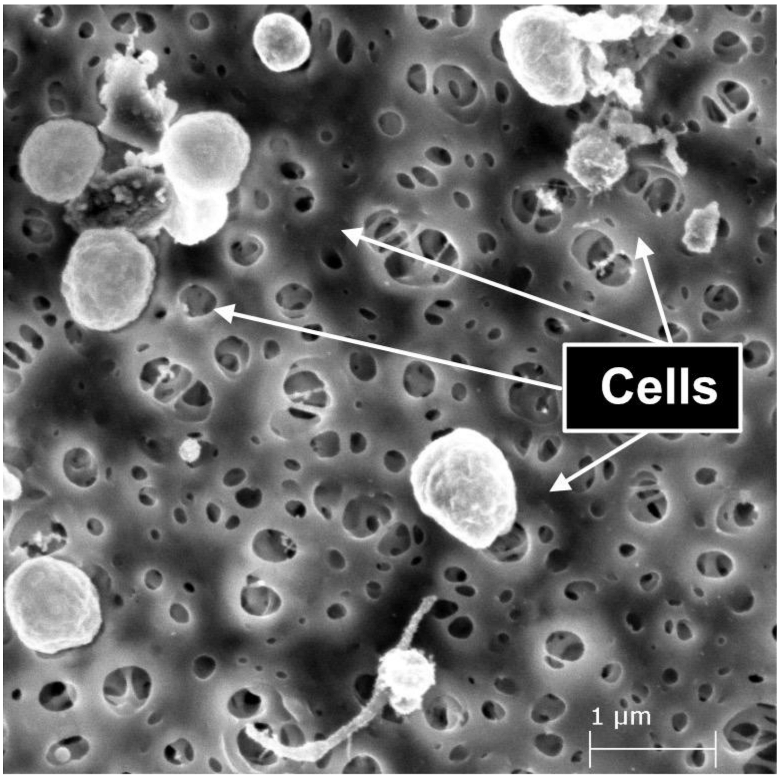


***Fig. S5.*** *Experimental control using M. bourgensis MAB1.*

***Table S3.*** *Raman shifts (cm$^{-1}$) of the spheres, and EPS.* The ($I_{1070}/I_{960}$) *is calculated on the phosphates and carbonates only (EPS do not represent Ca-P phases).*

| **Precipitates** | **Raman shift (cm$^{-1}$)** | **$I_{1069}/I_{960}$** |
|---|---|---|
| **Spheres 1/2** | 428/429 | 0.19 |
| | 588/585 | |

| | | |
|---|---|---|
| | 597/608 | |
| | 690/683 | |
| | 960/961 | |
| | 1072/1070 | |
| | 1305/1301 | |
| **EPS** | 953.5 | - |
| | 1064.0 | - |
| | 1129.0 | - |
| | 1235.0 | - |
| | 1296.5 | - |
| | 1442.5 | - |
| | 1632.5 | - |

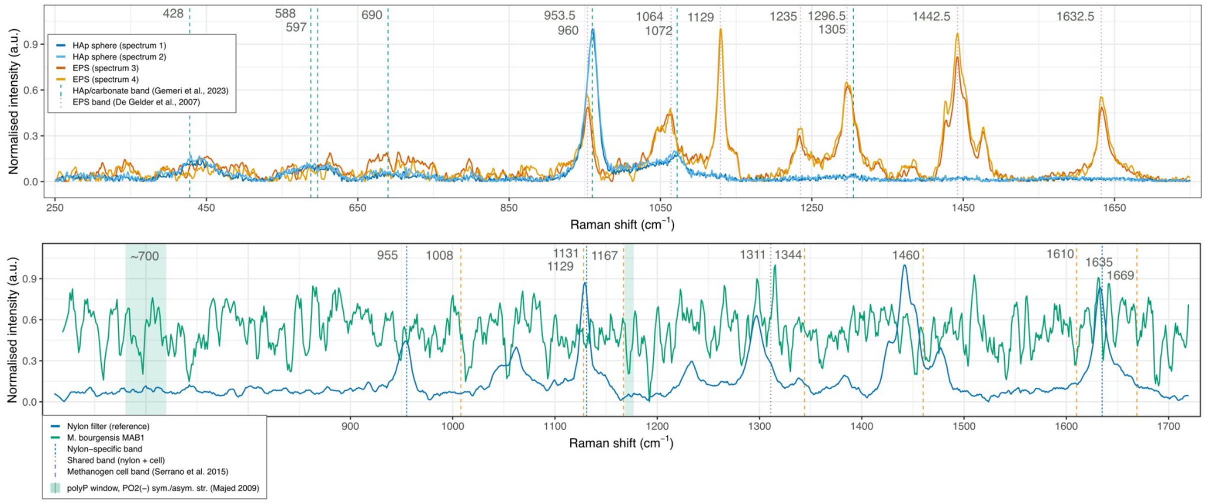


***Fig. S6.*** *Upper panel: Raman spectra of HAp spheres (blue) and EPS (orange and yellow). Vertical lines mark HAp and EPS specific bands (Gemeri et al. 2023; De Gelder et al. 2007). Lower panel: Raman spectrum of M. bourgensis MAB1 filament (green) and the nylon filter reference (blue). Vertical lines mark nylon-specific bands, bands shared between nylon and cells, and methanogen bands (*Serrano et al. 2015*). The shaded band marks the polyP diagnostic window (Majed et al. 2009).* No peaks resolve *above the noise floor at any of the* methanogen *cell bands or polyP positions.*

***Table S4.*** *pH values of first-batch experiments*

| [P] | pH (inoculated) | pH (blank) |
|---|---|---|
| **0.0** | 7.94 | 7.18 |
| **0.1** | 7.88 | 6.98 |
| **0.5** | 8.08 | 7.19 |
| **1.0** | 7.98 | 7.20 |
| **3.0** | 7.98 | 6.97 |
| **6.0** | 7.88 | 7.02 |
| **6.0 (M.oryzae)** | 7.98 | 6.98 |

| | | |
|---|---|---|
| **6.0 (MAB1)** | 7.85 | 7.12 |

***Table S5:*** *PhreeqC saturation modelling*

| Soln | Condition | pH | SI HAp | SI Calcite | SI Vivianite |
|---|---|---|---|---|---|
| 1 | P0 blank | 7.18 | 2.1 | 0.61 | -3.50 |
| 2 | P0.1 blank | 6.98 | 1.2 | 0.41 | -3.40 |
| 3 | P0.5 blank | 7.19 | 3.6 | 0.61 | -1.24 |
| 4 | P1 blank | 7.20 | 4.6 | 0.60 | -2.26 |
| 5 | P3 blank | 6.97 | 4.5 | 0.34 | -0.77 |
| 6 | P6 blank | 7.02 | 5.4 | 0.32 | -2.83 |
| 7 | P0 inoculated | 7.94 | 5.6 | 1.29 | -2.40 |
| 8 | P0.1 inoculated | 7.88 | 5.6 | 1.24 | -2.76 |
| 9 | P0.5 inoculated | 8.08 | 7.0 | 1.39 | -1.84 |
| 10 | P1 inoculated | 7.98 | 7.6 | 1.29 | -0.78 |
| 11 | P3 inoculated | 7.98 | 8.6 | 1.23 | -0.82 |
| 12 | P6 inoculated | 7.88 | 8.8 | 1.07 | -1.10 |

***Table S6a:*** *Prevalence of polyphosphate metabolism genes across 320 Methanobacteriota genomes*

| Gene | Genomes | Percentage | Function |
|---|---|---|---|
| *ppk1* (PF02503) | 77/320 | 24.1% | Polyphosphate synthesis (ATP → polyP) |
| *ppk2* (PF03976) | 49/320 | 15.3% | Alternative polyP synthesis (GTP/ATP → polyP) |
| *ppx* (PF02541) | 107/320 | 33.4% | Polyphosphate degradation (polyP → Pi) |
| **Complete pathway** | 79/320 | **24.7%** | Can both synthesise AND degrade polyP |

***Table S6b.*** *Distribution of polyphosphate genes across the 30 GTDB genera in the 320-genome tree.*

| Genus | n | *ppk1* | *ppk2* | *ppx* | Complete pathway (%) |
|---|---|---|---|---|---|
| Methanobacterium_F | 4 | 4 | 0 | 4 | 4 (100%) |
| JAUXXK01 | 1 | 1 | 0 | 1 | 1 (100%) |
| Methanarmilla | 8 | 7 | 0 | 7 | 7 (88%) |
| UBA349 | 6 | 5 | 0 | 5 | 5 (83%) |
| Methanobacterium_A | 19 | 14 | 0 | 14 | 14 (74%) |
| Methanobacterium_B | 30 | 15 | 1 | 17 | 15 (50%) |
| Methanobrevibacter_A | 4 | 2 | 0 | 2 | 2 (50%) |

| | | | | | |
|---|---|---|---|---|---|
| Methanobacterium_C | 4 | 1 | 0 | 1 | 1 (25%) |
| Methanosphaera | 23 | 5 | 0 | 17 | 5 (22%) |
| Methanobacterium (sensu stricto) | 27 | 5 | 0 | 5 | 5 (19%) |
| Methanocatella | 98 | 14 | 42 | 29 | 17 (17%) |
| Methanobacterium_D † | 23 | 3 | 0 | 4 | 3 (13%) |
| Methanobrevibacter | 25 | 0 | 6 | 1 | 0 (0%) |
| Methanothermobacter | 9 | 0 | 0 | 0 | 0 (0%) |
| Methanovirga | 7 | 0 | 0 | 0 | 0 (0%) |
| Methanobinarius | 6 | 1 | 0 | 0 | 0 (0%) |
| Methanothermobacter_A | 5 | 0 | 0 | 0 | 0 (0%) |
| JAHKLT01 | 3 | 0 | 0 | 0 | 0 (0%) |
| Methanorudis | 3 | 0 | 0 | 0 | 0 (0%) |
| UBA117 | 3 | 0 | 0 | 0 | 0 (0%) |
| Methanobaculum | 2 | 0 | 0 | 0 | 0 (0%) |
| Methanoflexus | 2 | 0 | 0 | 0 | 0 (0%) |
| CADBMS01 | 1 | 0 | 0 | 0 | 0 (0%) |
| CAZLGO01 | 1 | 0 | 0 | 0 | 0 (0%) |
| JBOFHT01 | 1 | 0 | 0 | 0 | 0 (0%) |
| Methanacia | 1 | 0 | 0 | 0 | 0 (0%) |
| Methanobacterium_E | 1 | 0 | 0 | 0 | 0 (0%) |
| Methanothermus | 1 | 0 | 0 | 0 | 0 (0%) |
| UBA587 | 1 | 0 | 0 | 0 | 0 (0%) |
| VGTM01 | 1 | 0 | 0 | 0 | 0 (0%) |
| **Total** | **320** | **77** | **49** | **107** | **79 (24.7%)** |

† Includes M. AMB_03, which places within Methanobacterium_D by tree topology. GTDB representatives alone: 2/22 (9%)

***Table S7***. *Comparison of AMB_03's ppk1 and ppx to complete-pathway genomes.*

| Genome | Genus | ***ppk1*** **identity (%)** | ***ppx*** **identity (%)** | ***ppk1*–ppx arrangement** |
|---|---|---|---|---|
| GCA_963787255.1 (AMB_03's sister) | Methanobacterium_D | 81.2 | 86.6 | same contig, same strand, ~280 bp apart |
| GCA_039666455.1 | Methanobacterium_D | 86.7 | 74.3 | adjacent, same strand |
| GCA_000012545.1 (*M. stadtmanae*) | Methanosphaera | 57.0 | 39.6 | adjacent, same strand |
| GCA_028890305.1 (*M. massiliense*) | Methanosphaera | 55.8 | 43.7 | adjacent, same strand |

***Table S8:*** *PLAAC Analysis Summary*

| Locus tag | Annotation | Putative functional relevance | Interpretation |
|---|---|---|---|
| M_AMB03_01659 | Peptidoglycan-binding domain protein | Cell envelope/surface interaction | Potential EPS or cell-surface scaffold |
| M_AMB03_00342 | Pseudomurein-binding repeat protein | Archaeal cell wall binding | Strong candidate for extracellular matrix association |
| M_AMB03_01390 | Ada metal-binding domain protein | Metal binding / DNA repair | Possible ion coordination / catalytic potential |
| M_AMB03_02928 | Glycosyltransferase family 39 protein | Polysaccharide synthesis | Possible role in EPS production |
| M_AMB03_00895 | Cupredoxin domain protein | Redox-active / metal-binding | Potential metal interaction at the extracellular interface |
| M_AMB03_02420 | Hypothetical protein | Unknown | Candidate amyloid-forming protein |
| M_AMB03_00929 | Hypothetical protein | Unknown | Candidate amyloid-forming protein |
| M_AMB03_00877 | Hypothetical protein | Unknown | Candidate amyloid-forming protein |
| M_AMB03_01937 | Hypothetical protein | Unknown | Candidate amyloid-forming protein |
| M_AMB03_00167 | Hypothetical protein | Unknown | Candidate amyloid-forming protein |
| M_AMB03_00026 | Hypothetical protein | Unknown | Candidate amyloid-forming protein |
| M_AMB03_02700 | Hypothetical protein | Unknown | Candidate amyloid-forming protein |

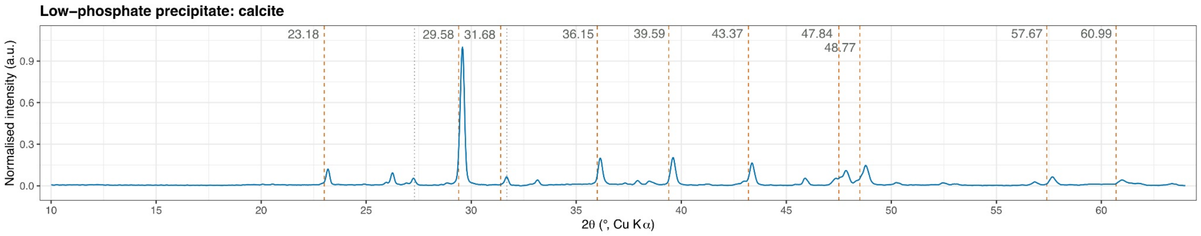


***Fig.S7.*** *XRD-pattern of the low-phosphate calcium carbonate precipitate, confirming calcite with minor halite. Data treatment (baseline subtraction,* Savitzky–Golay *smoothing) as described in 2.7.4. Kay et al. 1964; Graf 1961.*
